# Membrane curvature sensing and symmetry breaking of the M2 proton channel from Influenza A

**DOI:** 10.1101/2022.07.02.498578

**Authors:** Cole V. M. Helsell, Frank V. Marcoline, James Lincoff, Andrew M. Natale, Michael Grabe

**Affiliations:** Graduate Group in Biophysics, University of California, San Francisco, San Francisco, California, USA 93858-2330; Cardiovascular Research Institute, Department of Pharmaceutical Chemistry, University of California, San Francisco, San Francisco, California, USA 93858-2330

## Abstract

The M2 proton channel aids in the exit of mature influenza viral particles from the host plasma membrane through its ability to stabilize regions of high negative gaussian curvature (NGC) that occur at the neck of budding virions. The channels are homo-tetramers that contain a cytoplasm-facing amphipathic helix (AH) that is necessary and sufficient for NGC generation; however, constructs containing the transmembrane spanning helix, which facilitates tetramerization, exhibit enhanced curvature generation. Here we used all-atom molecular dynamics (MD) simulations to explore the conformational dynamics of M2 channels in lipid bilayers revealing that the AH is dynamic, quickly breaking the 4-fold symmetry observed in most structures. Next, we carried out MD simulations with the protein restrained in 4-fold and 2-fold symmetric conformations to determine the impact on the membrane shape. While each pattern was distinct, all configurations induced pronounced curvature in the outer leaflet with rather subtle lipid tilt, while conversely, the inner leaflets adjacent to the AHs showed minimal curvature and significant lipid tilt. The MD-generated profiles at the protein-membrane interface were then extracted and used as boundary conditions in a continuum elastic membrane model to calculate the membrane bending energy of each conformation embedded in different membrane surfaces characteristic of a budding virus. The calculations show that all three M2 conformations are stabilized in concave spherical caps and destabilized in convex spherical caps, the latter reminiscent of a budding virus. Only C2-broken symmetry conformations are stabilized in NGC surfaces, by 1-3 k_B_T depending on the AH domain arrangement. The most favored conformation is stabilized in saddles with curvatures corresponding to 33 nm radii. In total, our work provides atomistic insight into the curvature sensing capabilities of M2 channels and how enrichment in the nascent viral particle depends on protein shape and membrane geometry.

## Introduction

Influenza A M2 is a small, α-helical homo-tetrameric membrane protein essential for viral replication (Rossman & Lamb, 2011). M2 has two primary functions in the viral life cycle. First, during cell entry via endocytosis, endosomal acidification initiates fusion of the endosomal and viral membranes, but it also activates M2 channels causing further acidification that releases bound viral ribonucleoproteins (vRNP) to enter the host’s cytoplasm (Pielak & Chou, 2011). Second, during capsid exit from the cell, M2 channels residing in the host plasma membrane migrate to the budding virion where they enrich at the neck region and promote membrane scission resulting in the release of the mature membrane enveloped viral particle (Figure 1A). Consequently, M2-deletion mutants are replication impaired, resulting in accumulation of ∼25 nm blebs on the host membrane that only rarely resolve into infectious particles (Rossman et al., 2010). While other influenza factors are involved in viral egress, M2 seems to play the main part in ESCRT-independent scission of nascent viral buds by promoting further constriction of the existing ∼25 nm neck in order to facilitate spontaneous membrane scission, and thus determines the shape of the mature virion (i.e. spherical or filamentous) (Martyna et al., 2017; Rossman et al., 2012).

**Figure 1.**
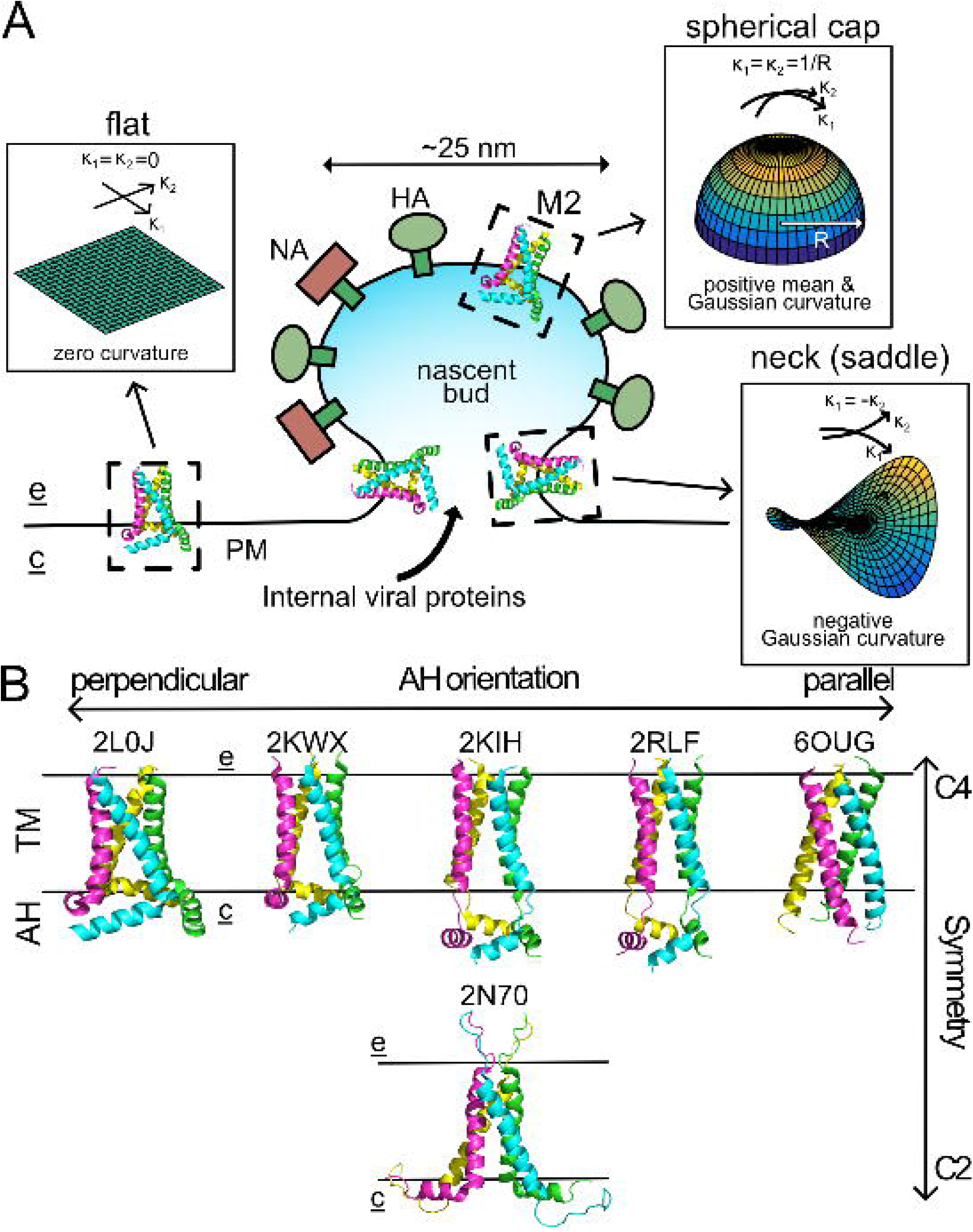
M2 channels and influenza egress. (A) Structural organization of the nascent viral bud. M2 first accumulates in the host plasma membrane, which is approximately flat and therefore has zero Gaussian curvature. M2 then migrates to the neck of nascent viral bud, which has negative Gaussian curvature. Finally, a small minority of the initial M2 population migrates into the approximately spherical cap of the mature virion. (B). A sample of published M2 structures (indicated by PDB accession codes) grouped by reported symmetry (top row: C4 symmetry vs. bottom row: C2 symmetry) and relative position of their amphipathic helix (AH) domains: 2L0J on the far left turns sharply and lies perpendicular to the TM domain, with the AH domain becoming less associated with the TM domain until 6OUG on the far right shows the AH continuing parallel to the TM helices.

The M2 protein is 97 residues long with largely disordered N- and C-termini connected by a single pass transmembrane α-helical segment (residues 23-46) and an amphipathic helix (residues 47-62) called the AH domain. Channels are homotetrameric assemblies with the ion conduction pathway formed at the center of the four transmembrane (TM) helices, and a minimal-construct “conductance domain” (CD) comprised of the TM and AH domains is both ion channel competent and sufficient for budding (Sharma et al., 2010a). The cytoplasm-facing AH domain is essential for scission activity (Roberts et al., 2013), but M2 chimeras with genetically unrelated amphipathic helices are scission-competent and promote viral replication *in vivo* (Hu et al., 2019). Meanwhile, the N- and C-termini are not essential for bud localization, NGC generation, or scission (Chen et al., 2008; Kwon & Hong, 2016; Martyna et al., 2017).

High-resolution structural insight comes from X-ray and solid state NMR (ssNMR) structures solved in a variety of conditions in the presence and absence of drugs, some of which are represented in Figure 1B. All structures reveal channels composed of four α-helical transmembrane segments, with varying degrees of transmembrane helix tilt with respect to the central axis. Generally, the helices exhibit greater splay on the cytoplasmic/intraviral side of the membrane than on the extracellular/extraviral surface giving rise to a conically shaped membrane spanning region. Helix kinking at glycine 34 has been suggested to control the degree of splay in active (greater) and inactive (less) conformations in a pH-dependent manner, which consequently increases and decreases, respectively, the conical character of the protein (Stouffer et al., 2008). Unfortunately, crystal structure constructs lack the entire AH domain and hence provide limited use in understanding the full membrane deforming function of the protein (Schmidt et al., 2013); however, several structures (mostly ssNMR) have resolved this portion revealing that it is the most variable part of the conductance domain (Figure 1B). In some structures it lies parallel to the membrane at the edge of the protein further exaggerating the conical shape of the channel (Andreas et al., 2015; Sharma et al., 2010b), but in others the elbow between the TM and AH domains is extended and the AH domains become more aligned with the membrane normal. In the extreme case, the partial AH domain of 6OUG forms a continuous helix with the TM domain. Most studies reveal that the backbone adopts a roughly 4-fold symmetric channel; however, a ssNMR structure of a seasonal variant of M2 provided spectroscopic evidence for a 2-fold symmetric tetramer (Andreas et al., 2015) in which the AH domains break 4-fold symmetry (Figure 1B). Moreover, double electron-electron resonance (DEER) experiments support the notion that the AH domains are dynamic with a high degree of flexibility (Herneisen et al., 2017; Kim et al., 2015).

Proteins induce membrane deformation via many mechanisms (Argudo et al., 2016; Farsad & Camilli, 2003; Jarsch et al., 2016; McMahon & Boucrot, 2015; Nepal et al., 2018) of which a subset have been proposed for M2 localization and curvature generation. Proteins that bind one membrane leaflet or transmembrane proteins with asymmetry across the bilayer midplane can impart their spontaneous curvature to the membrane via a *wedge insertion* mechanism, which has been proposed for a variety of membrane-bending proteins, such as epsins and BAR proteins. This mechanism is also relevant for M2 as the conically shaped CD construct is wedge-like, displacing more membrane area on the cytoplasmic leaflet than the extracellular leaflet, imparting a spontaneous curvature on the membrane that agrees with the estimated curvatures at the budding neck (Schmidt et al., 2013). An additional curvature-generating role for the AH domain is suggested by the *packing defect stabilization model* which states that bulky hydrophobic residues from amphipathic helices intercalate into the membrane at curvature-deformed sites to minimize exposure of the hydrophobic tails to solution (Cui et al., 2011). Finally, the nascent bud is cholesterol rich, giving rise to a line tension at the domain boundary with the host membrane. M2 and other membrane scission proteins have been shown to be “*linactants*“ that modify line tension (Kuzmin et al., 2005), supporting a role for M2 clustering at the boundary and aiding with demixing of the two phases (Madsen et al., 2018). A recent simulation study of M2 crowded into a bilayer patch showed the emergence of linear clusters and attendant changes in membrane curvature (Paulino et al., 2019). This *linactant model* provides the added benefit of explaining how influenza-specific membrane components are enriched from the host membrane (Gerl et al., 2012). A molecular mechanism supporting M2’s role as a linactant comes from experiments showing that M2 binds cholesterol sub-stoichiometrically with two proximal (as opposed to diagonal) lipids per M2 tetramer (Elkins et al., 2017). This asymmetry allows M2 to satisfy its preferred cholesterol binding orientation when it localizes to the interface and one side embeds in the cholesterol rich bud and the other faces the host membrane.

While these theories of curvature generation suggest that M2 may deform membranes, they do not address the observation that M2 migrates to the neck of budding vesicles as opposed to the spherical cap, for instance. In fact, the dominant view of M2 as a 4-fold symmetric, wedge-like shape is at odds with NGC localization, because any two orthogonal axes of the protein induce spontaneous curvatures of the *same* sign (Figure 1A). However, the neck is a catenoid or saddle-like membrane structure characterized by two, orthogonal principal curvatures, κ_1_ and κ_2_, of *opposite* sign (Figure 1A). Moving along either principal direction, the membrane in the neck bends away from a point on the surface in different directions, and hence the Gaussian curvature, defined as K = κ_1_·κ_2_, is negative. Thus, while the shape of M2 may curvature-match the neck in one direction, it would appear to frustrate the local shape in the orthogonal direction suggesting that, if anything, the channel should be excluded from the neck. Recent coarse-grained simulations by Voth and coworkers using a 4-fold symmetric model of M2 support this notion, as they found that M2 migrated to patches of positive gaussian curvature rather than the expected NGC regions (Madsen et al., 2018).

One solution to this apparent contradiction is that M2 and the AHs break 4-fold symmetry, as discussed above, and adopt configurations compatible with the two distinct principal curvatures present in the neck. Tangential support for such a mechanism comes from work by the Antonny lab showing that geometric rearrangements of defect-sensing helices imparted by changes in the flanking protein scaffold have modified curvature specificity in the cell (Doucet et al., 2015). Here, we initiated unrestrained, all-atom MD simulations of the M2 channel and identified large conformational changes in the AH domain that broke 4-fold symmetry. Inspired by these broken symmetry conformations, we next asked how the surrounding lipid bilayer responded to M2 channels with different symmetry configurations. To do this we performed restrained simulations on 3 different systems: a 4-fold symmetric NMR structure, a 2-fold symmetric NMR structure, and a 2-fold symmetric parallel AH domain model inspired by our unrestrained simulations. Each protein configuration induced specific deformations in the surrounding bilayer with 2-fold distortion patterns coming from the latter two systems and a largely radially symmetric distortion pattern emitted from the 4-fold channel structure. However, all systems shared several common features, notably pronounced curvature generation and correspondingly very little lipid tilt in the extracellular leaflet and a mostly flat cytoplasmic leaflet exhibiting strong lipid tilt. Using the MD-derived deformations at the channel boundary as boundary conditions, we determined the membrane energetic cost associated with placing these 3 different protein configurations into spherical caps, flat membranes, and saddle regions. None of these configurations are predicted to be stabilized in the convex, positive mean curvature spherical cap of a budding vesicle, while they all favor concave, negative mean curvature spherical caps. The two configurations exhibiting 2-fold symmetry are stabilized in negative Gaussian curvature saddles by 1-3 k_B_T over a flat membrane, while the 4-fold symmetric structure is not. Additionally, the parallel AH domain structure favors a saddle with a radius of curvature of ∼30 nm very close in size to the stalled particle size observed in cells infected by M2 defect mutant strains (Rossman et al., 2010). Thus, our results provide a mechanical mechanism for why M2 channels are absent in the spherical cap of a nascent capsid and suggest that symmetry breaking by the amphipathic helices could explain channel localization to the neck region.

## Results

### Unbiased MD simulations reveal flexible amphipathic helical regions

To explore the conformational landscape of the M2 protein, we initiated an unbiased all-atom MD simulation of the 4-fold structure (PDBID: 2L0J) in a heterogeneous bilayer consisting of POPC, POPG, and cholesterol. Specifically, the backbone of 2L0J exhibits 4-fold symmetry providing the impression that it is globally symmetric, yet the individual side chains on each monomer lack any symmetry, while several histidines in the central pore (not in contact with the membrane) were built with 2-fold symmetry. Just after equilibration at the start of production, the channel (AH - red/TM - yellow) was quite similar to the starting NMR structure (transparent white) (snapshot 1, Figure 2A). However, over the first 500 ns, the total protein RMSD (blue curve) increased, and the AH domains (red curve) mirrored this increase but were even larger by 1-2 Å reaching just over 7 Å at 500 ns (snapshot 2). Several AH helices have broken symmetry and moved radially away from the central axis losing contact with the other subunits, with the top helix the most pronounced in this example. In contrast, the TM domain (yellow) remained stable throughout the 2,750 ns simulation with an RMSD ∼1 Å. Additional snapshots along the trajectory (3-5) reinforce the notion that the AH helices are mobile, showing that they all deviate from the starting structure, some quite significantly. The helices primarily retain their helical secondary structure, but some partially unfold and sometimes refold (top helix in snapshot 4 versus 5). The sixth and final snapshot shows a side view of the channel highlighting that the TM domains (yellow) retained 4-fold symmetry with a very close match to the 2L0J structure.

**Figure 2.**
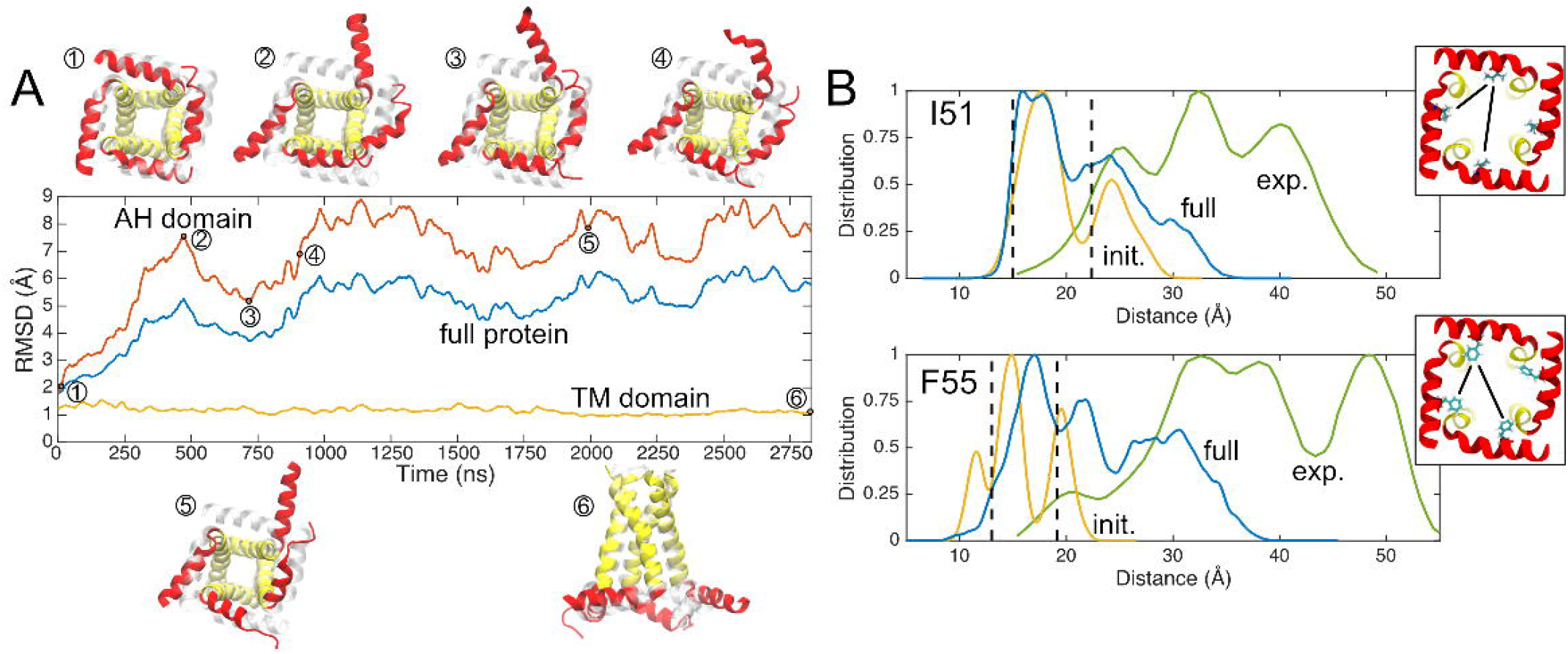
Unrestrained simulations reveal dynamic AH domains. (A) RMSDs of Ca backbones over time. Yellow indicates the TM domain only, red the AH domain only, and blue the full protein. Snapshots from unrestrained MD are numbered along the X axis with the starting structure overlaid in light grey, showing AH dynamics. (B) DEER data provided by the Howard lab (green) compared to distance estimates from simulation between labelled AH domain residues (blue), including the first five nanoseconds alone (yellow). Insets in upper and lower panels show the I51 (top) and F55 (bottom) residue on 2L0J (view from cytoplasm), with adjacent and diagonal distances represented by two black lines.

We computed the residue-to-residue distance distribution for two AH residues (I51 and F55), which had previously been measured using DEER for the same construct in the same lipid composition (Kim et al., 2015). Each residue has essentially two distal heavy atom distance values in the 4-fold starting structure, a smaller adjacent subunit-to-subunit value (∼13 Å) and a larger diagonal value (18 or 22 Å) indicated by vertical dashed lines (Figure 2B). The experimental distances (green) are much broader and occur over larger distances ranging from 20-45 Å for the I51 position and 20-50 Å for the F55 position. Part of this difference arises from the increased length of the MTSL probes compared to the isoleucine and phenylalanine side chain surrogates measured from the structure, but this difference alone cannot explain the large discrepancy. The dynamic nature of the AH helices provides a picture much closer to the experimental values, as proposed by the Howard lab (Kim et al., 2015). We computed the inter-subunit amino acid-to-amino acid distance distributions for both residues (6 unique distances for each position in the 4-subunit channel) from the trajectory over very short times 5 ns (yellow curve) and the full 2,750 ns simulation (blue curve). Initially the side chains explore a range of rotamer conformations, despite little backbone motion in the AH domain, and this results in distributions that are several ångströms wider than the bounds extracted from the NMR structure, but fail to account for the DEER data. As the simulations progress, the deviations in the AH domains and breaking of 4-fold symmetry create a much broader range of distance values (blue curves) 15-20 Å greater than the largest values extracted from the static structure that begin to approach the experimental range. The lack of agreement between simulations and experiment is not unexpected as we are not modeling the probes explicitly, and our simulations are not converged. This latter point is evident as we see excursions in some AH helices that are not sampled in others despite the symmetry.

Our unbiased simulations confirm that the AH domains are flexible and can adopt many conformations, some of which break the 4-fold symmetry of the channel. Both of these observations are consistent with the range of conformations observed in structural databases (see Figure 1) and with DEER experiments (Herneisen et al., 2017; Kim et al., 2015). The simulations also suggest that reorientation of the AH domains in a flat bilayer requires little energy and may be favorable, as it occurs spontaneously on a short timescale (microseconds) and in some cases helices return to their starting position (right AH helix in snapshot 2 versus 3, Figure 2A).

### M2 channels deform the membrane

Next, we wanted to quantitatively investigate the characteristics of the membrane around the M2 channel and explore how its properties depend on the conformation of the protein. Specifically, we were interested in the role that C4 and C2 symmetric channels play in patterning the local membrane shape, and we probed this question by carrying out protein-restrained, all-atom MD simulations on three different conformations: 1) the 4-fold 2L0J structure (Sharma et al., 2010b), 2) the 2-fold 2N70 structure from the Griffin lab (Andreas et al., 2015), and 3) a 2-fold symmetric parallel AH domain model based on conformational changes observed in our unbiased simulations of 2L0J (Figure 3A). By restraining the backbone conformations to the starting structures, we could explore how the membrane relaxed around specific channel conformations in an attempt to explicitly link protein shape to membrane deformation. The parallel AH domain structure was inspired by the anecdotal observation in our unrestrained 2L0J simulation that AH domain helices from adjacent subunits break symmetry and point parallel to each other in the plane of the membrane (snapshots 3 and 5 in Figure 2A). We expected that such an arrangement would also break symmetry in its patterning of the membrane, potentially inducing different curvatures and hence influencing the curvature sensing of the channel. To create thisMmodel, we started from the 2N70 structure, removed the unfolded AH domains, and replaced them with one of the folded AH domains from the adjacent subunits. The two helices are largely parallel to each other, similar to the top and right helices in snapshot 5 shown in Figure 2A, with a slight rotation along the AH helical axis.

**Figure 3.**
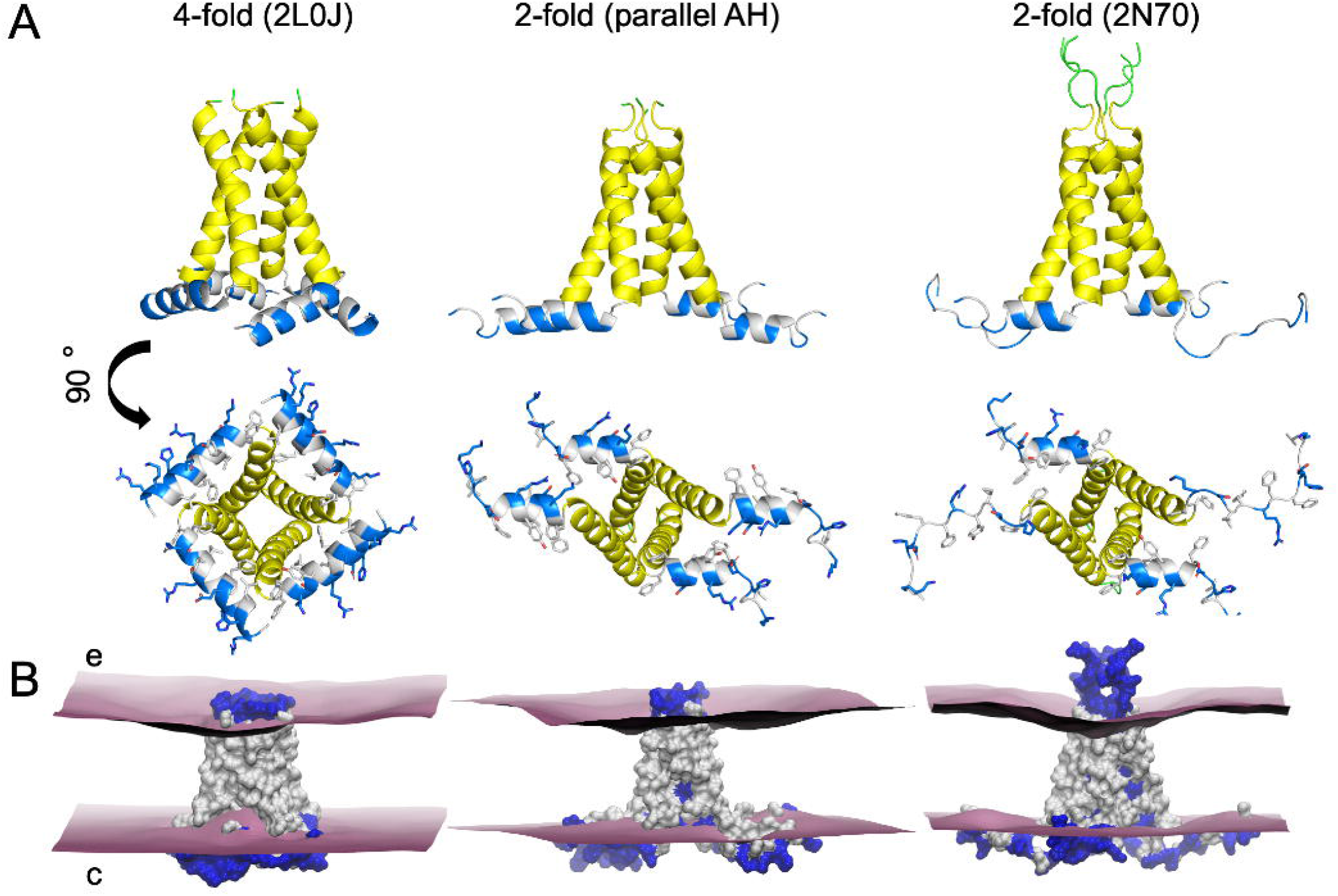
M2 membrane deformation patterns from simulation. (A) Structures for restrained-protein simulations. Extracellular loops in green, TM domain in yellow, AH domain in blue (polar/charged) and white (hydrophobic). Left: the 4-fold solid-state NMR structure (PDBID 2L0J); center: parallel AH domain model (see main text for details); right: 2-fold solid-state NMR structure (PDB ID 2N70). (B) Upper and lower mean membrane surfaces computed from MD (purple). Polar and charged sidechains shown in blue, hydrophobic sidechains in white.

The organization of each channel is pictured in Figure 3A shown from both the side (membrane view) and cytoplasm. The cytoplasmic view highlights the distinct symmetry of each configuration, and while the parallel AH domain model and 2N70 are C2 symmetric, their overall configurations are quite distinct due in large part to the extended AH domains of 2N70. The AH domain is amphipathic consisting of charged and polar (both in blue) and hydrophobic residues (white) with the side chains explicitly represented in each structure. As expected for an amphipathic helix, the charged/polar residues point down into solution and the hydrophobic residues point up into membrane core. This is also true for the amino acids on the extended AH domains of 2N70 as the phenylalanines (white) and acidic/basic (blue) residues extend their side chains in opposite directions.

Each restrained simulation lasted between 1.7 to 3.8 μs (Table 1), and the membrane deformation pattern induced by the proteins relaxed to an equilibrium shape after several hundred nanoseconds, except in the case of 2L0J which exhibited an upward movement of the lower leaflet by several *å*ngströms around 1 μs before becoming more stable from 1 to 3.8 μs (Fig. S1). Thus, we report all membrane properties averaged over the second half of each trajectory. The mean upper and lower surfaces representing the interface between the headgroups and the lipid tails is depicted around each protein, which is shown in surface view with hydrophobic residues white and charged and polar residues blue (Figure 3B). The charged portions of the AH domains fall primarily below the lower surface indicating that they are embedded in the headgroup regions or directly exposed to solution. Meanwhile, the residues exposed to the membrane core between the upper and lower leaflets are almost entirely hydrophobic. In all three cases, the membrane pinches, or compresses, as it approaches the channel, and this feature is accompanied by a pronounced curvature in the extracellular leaflet, while the cytoplasmic leaflet remains rather flat. Despite the presence of conserved general features, the deformation patterns are different.

**Table 1.** List of simulations.

| ID | Gromacs | Atoms | Length (ns) | PDBID | Bilayer composition | Restraints |
| --- | --- | --- | --- | --- | --- | --- |
| 1 | 2020.6 | 96,411 | 2,829 | 2L0J | 196xPOPC, 49xPOPG, 105xCHOL | No |
| 2 | 2020.6 | 96,351 | 3,808 | 2L0J | 196xPOPC, 49xPOPG, 105xCHOL | Yes |
| 3 | 2020.6 | 97,591 | 3,760 | parallel AH | 196xPOPC, 49xPOPG, 105xCHOL | Yes |
| 4 | 2020.6 | 117,473 | 1,727 | 2N70 | 196xPOPC, 49xPOPG, 105xCHOL | Yes |

Heat maps of the upper (top row) and lower (middle row) membrane surfaces from Figure 3B provide a more detailed view of the distortions (Figure *4*). The X and Y axes correspond to the size of one periodic image of the simulation box, and the inner black curves represent the shape of the membrane-protein contact curve in the upper and lower leaflets. The footprint of each channel is much larger in the lower leaflet than the upper leaflet, resulting from the presence of the AH domains and providing M2 its characteristic wedge shape. Additionally, the 4-fold versus 2-fold symmetry of each conformation is apparent in the contact curves. Membrane-thinning deflections of the surface, downward for upper leaflet and upward for the lower leaflet, are signed as negative, and outward deflections (membrane thickening) are positive. The mean nominal undistorted height of each leaflet was set to zero in each surface based on the membrane heights at the edges of the simulation boxes.

**Figure 4.**
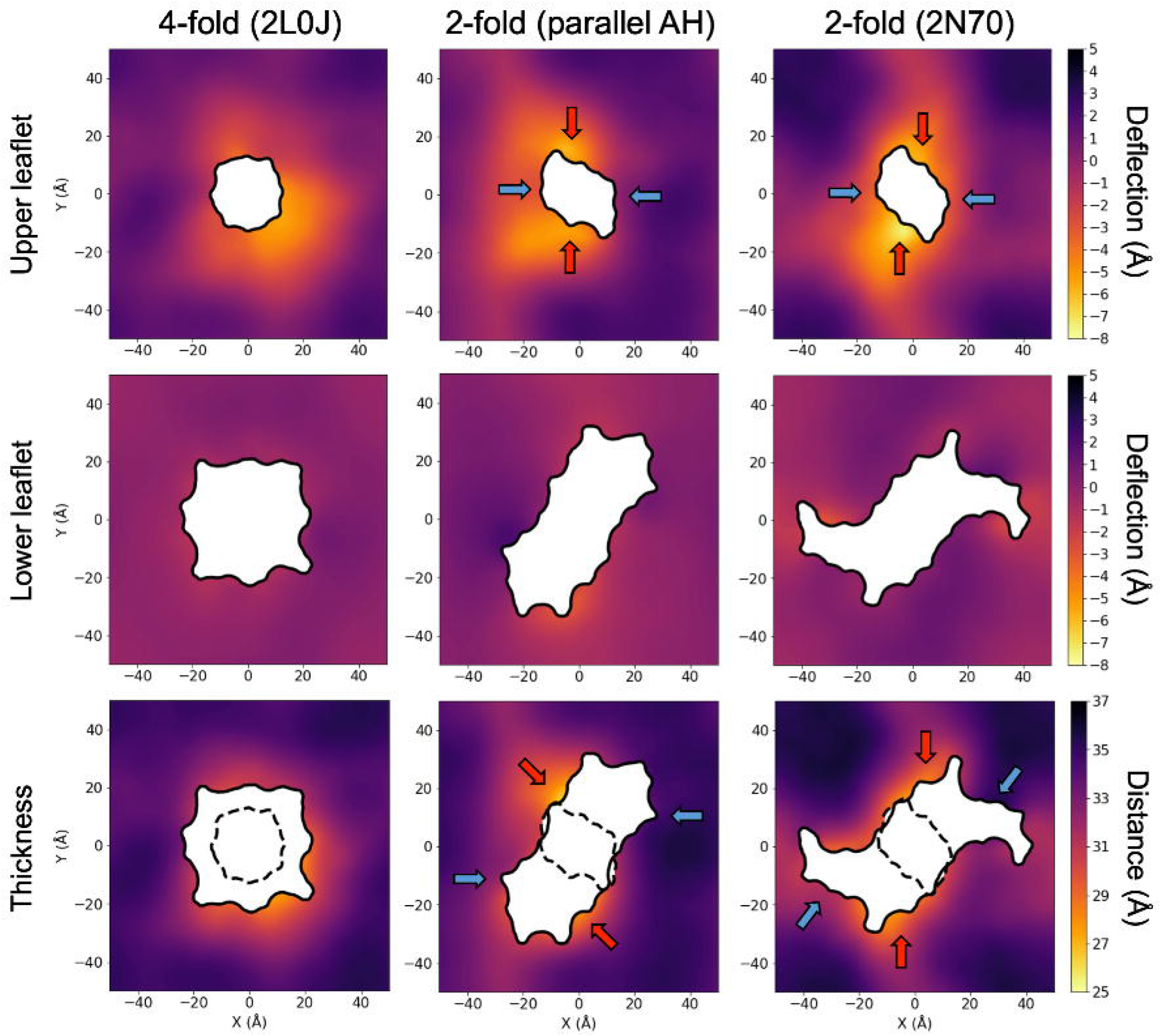
Surface height and thickness heatmaps for three different channel configurations. Compressive deflections are negative/shown in bright colors. Expansive deflections are positive/dark colors. The same scale is used for the upper and lower leaflets to highlight the greater amount of deflection in the upper as compared to the lower leaflet. Arrows correspond to regions of high (red) and low (blue) compression noted in the text.

The membrane distortion for 2L0J (left column) shows a high degree of rotational symmetry consistent with its C4 symmetry. At the membrane-protein contact curve, the upper surface is deflected downward by -3 to -5 Å while the lower leaflet surface is almost completely flat throughout. Moving radially away from the membrane-protein contact curve, the upper deflection returns to its equilibrium value with 0 Å deflection about 12-15 Å away from the protein. Thus, the upper leaflet exhibits membrane-bending curvature, while the thickness of the membrane (bottom row) reveals a pinch at the point of contact with the protein by 3-5 Å, indicating that the membrane is also under compression. In our simulations, far from the proteins the membrane adopts an equilibrium thickness of ∼35 Å consistent with experimental work suggesting PC:PG:cholesterol bilayers are ∼33 Å thick (Tong et al., 2012). The degree of pinching will likely vary for different lipid compositions.

Remarkably, the lower leaflet is flat in all three simulations with almost no sign of induced curvature despite the presence of the partially inserted AH domain helices. Moreover, the AH helices from each conformation are positioned differently in the leaflet with 2N70 only having two helical AH domains while the other two are unraveled, yet the lower leaflet membrane behaves similarly in each case. Like the 4-fold 2L0J system, the 2-fold structures induce bending and pinching in the upper leaflet, but to an even greater degree. Contrary to the 2L0J results, the parallel AH domain model (middle column) and the 2N70 structure (right column) both break membrane symmetry with the largest negative deflection happening in the upper leaflets along the Y-axes and less negative deflections along the X-axes (red and blue arrows in top row, respectively). The parallel AH domain model induces a -5 to -7 Å deflection at these points of contact between the protein and membrane, but only 0 to -3 Å deflection along the X-axis contact points. The upper leaflet in the parallel AH domain model generally shows greater negative deflection for the X < 0 region of the plane than the X > 0 side. The 2N70 structure sets up a trough-like distortion in the upper leaflet along the Y-axis with large negative deflections along the ± Y-axis contact points (−7 to -8 Å) and modest defections along the ± X-axis contact points (∼ -3 Å). Taking the upper and lower deflections together, the membrane is pinched by 5 Å at the point of contact with the protein along the -Y axis and by about 4 Å at the point of contact along the +Y axis (red arrows in Figure *4*), while along the X = Y diagonal it shows two symmetric regions of very little compression (blue arrows). The resulting compression pattern for 2N70 is somewhat similar to the parallel AH domain model with symmetric regions of compression (red arrows) and symmetric regions of modest to no compression (blue arrows). The different channel conformations thus produce distinct local membrane deformation patterns that reflect their individual symmetries and AH orientations, suggesting that such conformational changes may be key to the dependence of local M2 enrichment on membrane geometry seen in experiment (Rossman et al., 2010).

### Lipids tilt around the amphipathic helices

We next characterized the lipid tilt in the three restrained simulations, beginning with a qualitative assessment on individual snapshots of the side/membrane view of the proteins in equilibrated membranes (Figure *5*, top row). While the bilayers contain POPC, POPG, and cholesterol, cholesterol was excluded in the tilt calculations. The proteins here are oriented to match the side views in Figure 3, with the viewpoint origin in the lower right quadrant of the tilt surfaces (Figure *5*, lower rows). First, the upper leaflet curvature discussed in the previous section and Figure *4* is visible in these snapshots for 2L0J (left column) and 2N70 (right column) though not the 2-fold parallel AH domain model (middle column), and all lower leaflet surfaces appear comparatively flat. However, there is far greater lipid tilt apparent in the lower leaflet, arising from a mixture of lipid tail kinking (e.g., the purple POPC in the 2N70 snapshot), splaying apart of lipid tails (e.g., the purple POPC in the parallel AH domain model snapshot), and simple off-normal tilting of an otherwise “regular conformation” lipid (e.g., the purple POPG in the 2L0J snapshot). These various tilt behaviors appear most pronounced near the proteins, as lipids reorient to conform to the irregular wedge-shape of the channel created by the partial insertion of the AH domains in the lower leaflet, while more distant lipid solvation shells exhibit less pronounced tilt. Small amounts of tilt can be seen in the upper leaflet as lipids orient around the tilted TM helices (most strongly for 2N70), though they retain largely normal conformations typical of lipids in a flat, protein-free bilayer. Of note, while we observe cholesterol adjacent to the channel, and cholesterol has been shown to bind M2 (Elkins et al., 2017), our simulations failed to identify any binding events or hot spots on the low microsecond timescale, which may be the result of the backbone restraints or inadequate sampling.

**Figure 5.**
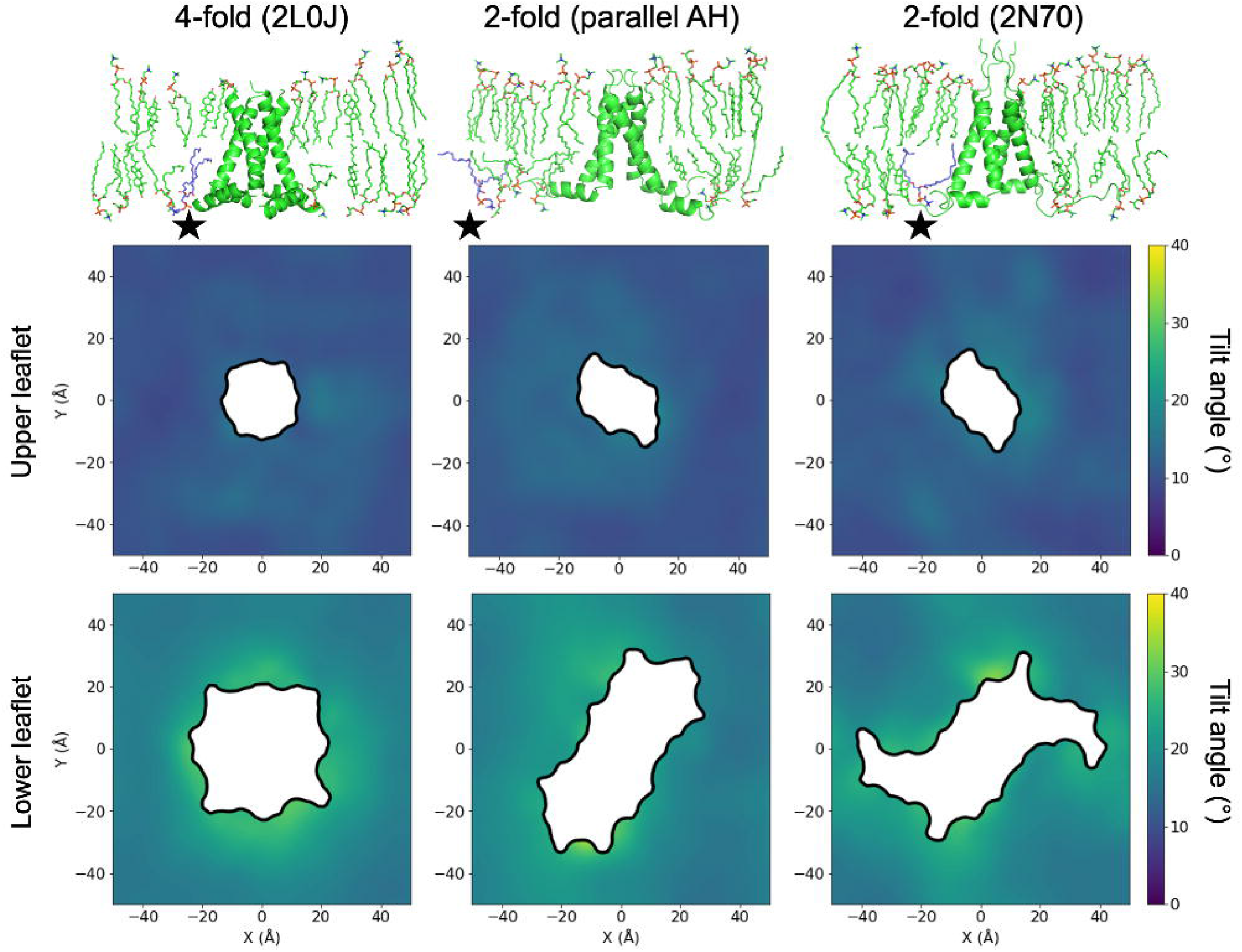
Lipid tilt around different M2 conformations. (A) Representative all-atom snapshots extracted from the protein-restrained, equilibrium simulations of 2L0J, parallel AH domain model, and 2N70 (2,3 and 4 in Table 1, respectively). Black stars in each snapshot highlight purple lipids discussed in the main text. (B) Two-dimensional tilt surfaces computed for each simulation in panel A separated out by upper (middle row) and lower (bottom row) leaflets. The mean lipid tilt at a given position in the x-y plane with respect to the z-axis (membrane normal) is reported in degrees and color coded according to the scale on the right. Middle and bottom rows correspond to simulation system in top row.

Lipid tilts throughout the membrane surface were then quantified in the same way as the leaflet deflections and membrane thicknesses were (Figure *5*, middle and bottom rows), with the individual phospholipid tilts calculated as the angle between the presumed membrane normal (aligned with the Z-axis of the box) and the vector pointing from each phospholipid’s phosphate to the midpoint between the last carbon atoms of the lipid tails. For all conformations, the mean tilt is consistently higher in the lower leaflet (bottom row) than the upper leaflet (middle row), as evidenced by the brighter colors of the lower leaflet surfaces than the upper leaflets. For the upper leaflet, there are spots of higher tilt (∼20°) at the protein interface for 2N70, but otherwise the tilt in the upper leaflet for all simulations is bounded between 10° and 15°. The universally greater degree of tilt in the lower leaflet is consistent with the hypothesized impact on adjacent lipids of AH partial insertion studied by May and co-workers (Zemel et al., 2008), reviewed by Zimmerberg and Kozlov (Zimmerberg & Kozlov, 2006), and recently observed with brominated lipid probes and coarse-grained MD in an ESCRT-III system (Moss et al., 2021). It is also possible that the greater splay of the TM helices on the cytosolic leaflet also contributed to the increased lipid tilt. In all simulations, lower leaflet tilt is highest—around 30 to 35—at the membrane-protein contact curve, and then decays to lower values of around 15° toward the edges of the simulation cell. The tilt pattern in the lower leaflet for 2L0J seems nearly rotationally symmetric, as might be expected from the 4-fold symmetry of the construct. The 2-fold symmetric constructs produce less-rotationally symmetric tilt patterns.

To further quantify the patterning of the tilt around the proteins, we plotted the values of the mean tilts in the leaflets that make up the surfaces in Figure *5* against the distance from the protein surface (Figure *6*). Each grid point is then colored according to the local density of points around them, such that bright spots indicate regions where there is a high density of points that sample similar values of mean tilt versus distance from protein. Scanning vertically through the swath of points at a fixed value of distance from the protein thus highlights characteristics of the tilt distribution at a certain distance from the protein. The range that is sampled reflects whether the distribution is narrow or broad, and the number of brighter spots seen at a fixed distance reflect whether there is a single common mean tilt or multiple frequently-sampled mean tilts at the same distance from the protein.

**Figure 6.**
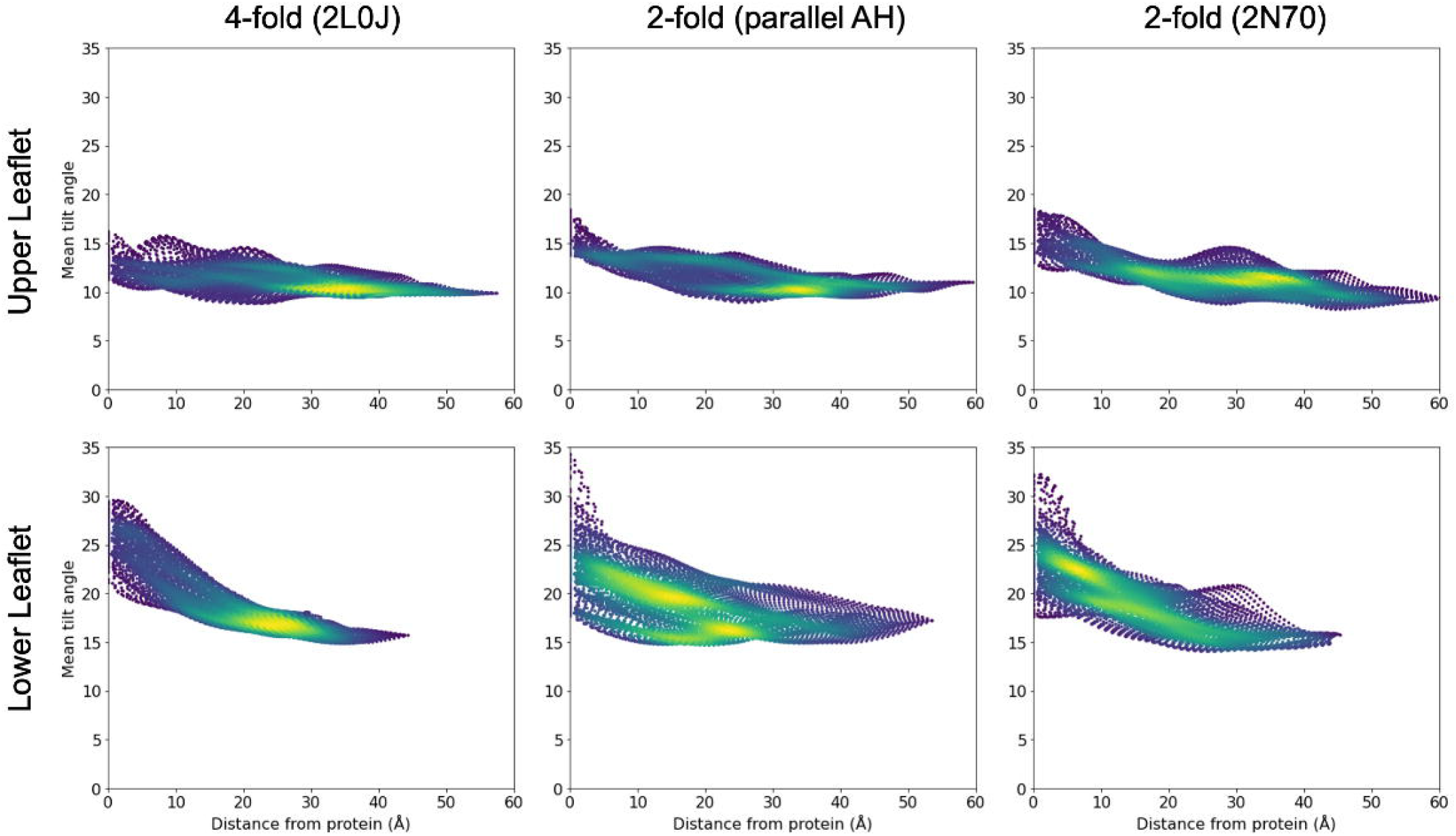
Mean tilt as a function of distance from protein for restrained simulations. Points are colored by local density, such that bright spots reflect highly-sampled values of tilt versus distance. At a fixed value of distance, a vertical scan highlights the range of mean tilts sampled and the nature of the distribution of mean tilts seen at that distance.

As expected from the tilt surfaces in Figure *5*, the 2L0J simulation (left column) produces the most isotropic patterning of tilt versus distance of the three simulations. In the upper leaflet, the tilt decreases from a mean value of 12° at the protein to 10° at 30 Å and beyond. The lower leaflet displays a much stronger decay of tilt, with an apparent uniform distribution of 21 to 30° near the protein that decreases to a mean around 17° by around 20 Å of distance. The ranges of tilts seen in the 2-fold symmetric simulations largely match those seen for 2L0J, but the distributions are rather different. The 2-fold parallel AH domain structure (middle column) shows multiple tilt modes in the upper leaflet at a distance of 20 Å from the protein, at 10 and 14°. Multiple common tilt modes are also present in the lower leaflet immediately around the protein, highlighting the X > 0 versus X < 0 anisotropy seen in Figure *5* for the parallel AH model. The 2N70 simulation shows somewhat less anisotropy, with both leaflets dominated by a single mean tilt value at all distances from the protein, except values less than 10 Å where bright spots are present around 12 and 16° in the upper leaflet and around 19 and 23° in the lower leaflet. Overall, though these simulations produce similar ranges of tilt for the upper and lower leaflets, the appearance of different multi-modal tilt distributions at various distances from the protein highlights the strong influence that the protein conformation has on the local tilt field, which may factor into the aggregation and localization behavior of the protein.

Contrasting the height deformation profiles in Figure *4* with the tilt profiles in Figure *5*, a striking feature emerges: all M2 conformations induce large curvature in the upper leaflet with hardly any curvature in the lower leaflet, while the lipid tilt profiles experiences very little deviation from a flat bilayer in the upper leaflet and strong deviation in the lower leaflet. At the atomic level, the snapshots in Figure *5* explain how the different M2 channels confer this behavior. The lower leaflet lipids adjacent to the AH domains kink and tilt their tails to wrap around the helices. These deformations reduce the vertical extent that the tails can reach across the membrane, and the upper leaflet lipids at the protein interface move down to fill any gaps, which produces the pinching observed in the outer leaflet. Meanwhile, the extracellular span of M2 presents a cylinder-like surface to the surrounding lipids compatible with small lipid tilt angles aligned along the membrane normal. Retaining small, bulk-like lipid tilt angles in the upper leaflet also makes it possible for these lipid molecules to extend farther across the bilayer further compensating for the reduced reach of the lower leaflet lipids.

### A continuum membrane model

Next, we calculated the membrane deformation energy associated with each of these three M2 configurations to determine whether they would preferentially migrate to cellular regions of specific curvature such as the flat, saddle, or spherical caps relevant to a nascent viral bud (Figure 1A). Unfortunately, membrane deformation energies cannot be directly determined from our atomic simulations without employing sophisticated free-energy calculations (Fiorin et al., 2020), and there is no atomistic framework for determining the relative energetics of these membrane proteins in different background curvature fields, as the periodic boundaries in all-atom MD are incompatible with simple saddle shapes and spherical caps. Thus, we decided to apply continuum membrane mechanics to answer these questions. Previously, we developed a hybrid atomistic-continuum approach for determining the insertion energies and induced membrane distortions of integral proteins (Choe et al., 2008) that we extended to accurately represent the membrane around proteins of complex shape (Argudo et al., 2017). The membrane model is based on a Helfrich Hamiltonian (Helfrich, 1973) to account for elastic curvature in each leaflet coupled with a “mattress model” accounting for compression between the leaflets (Huang, 1986):

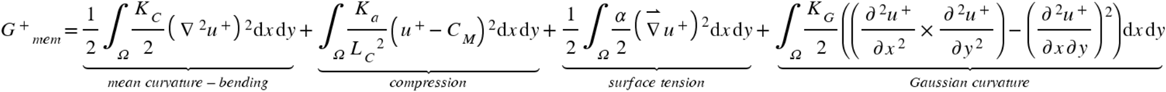

where for brevity we have only written the energy terms for the upper leaflet, u^+^ is the deviation from a flat equilibrium height for the upper leaflet, K_C_ is the mean bending modulus, K_a_ is the areal compression modulus, K_G_ is the Gaussian bending modulus, α is the surface tension, C_M_ is the bilayer mid-plane, and the integrals are carried out over the entire X,Y extent of the membrane domain Ω. Minimizing the full free energy for both leaflets, G_mem_, results in a set of Euler-Lagrange equations that are solved to determine the membrane distortions and subsequently the membrane distortion energy (Argudo et al., 2017). These solutions require numerically solving a 4^th^ order boundary value problem that involve setting the displacement and slope of the membrane at the protein-membrane contact curve (Figure *7*A) and on the far field boundaries. Previously, we applied a similar membrane model in an attempt to predict the protein-membrane interface for M2 channels lacking the AH domains (Zhou et al., 2020); however, here we directly used the MD generated membrane distortions at the protein boundaries to parametrized this boundary value problem.

**Figure 7.**
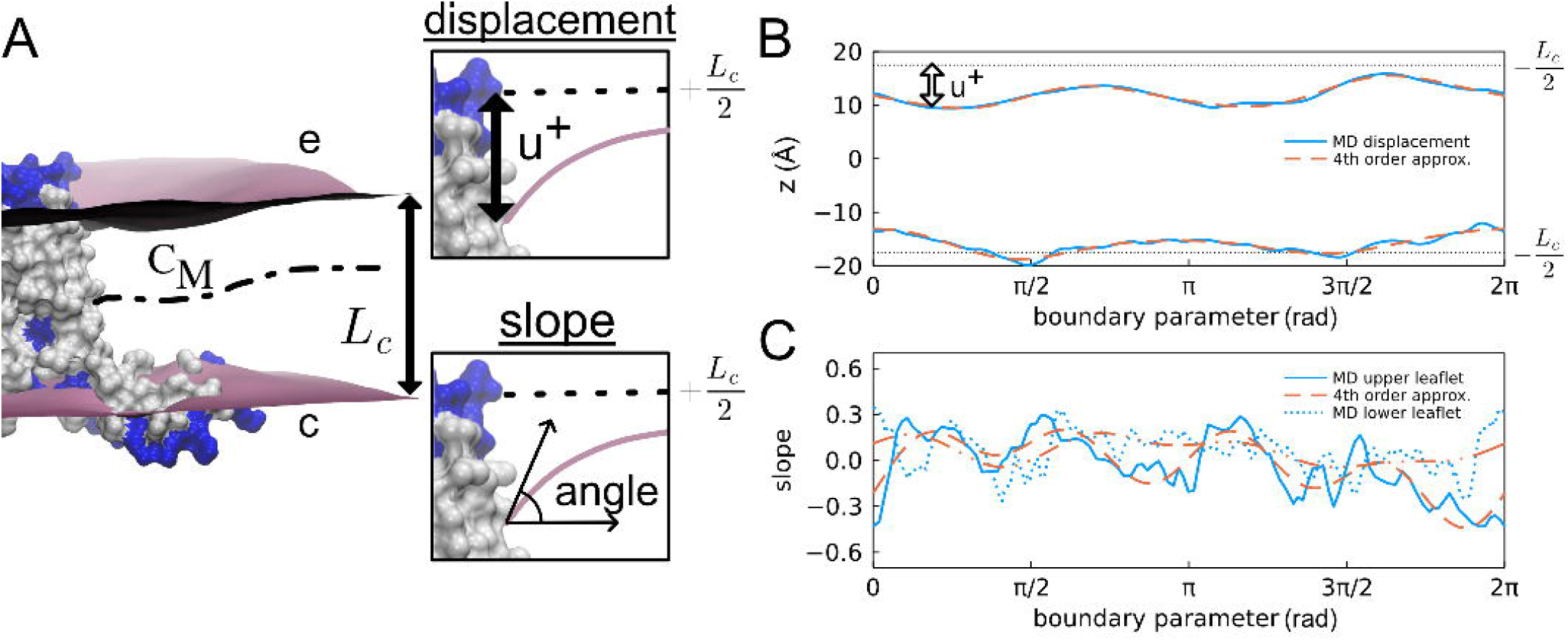
Boundary conditions extracted from MD simulations. (A) Parallel AH domain M2 protein in a lipid bilayer. The molecular surface is shown with hydrophobic residues in white and charged and polar residues in blue. The mean hydrophobic core of membrane, with equilibrium thickness L_c_, is between the purple surfaces. The mid-plane surface between the upper and lower leaflets is C_M_. The insets show the boundary conditions for the upper leaflet extracted at the lipid excluded surface of the protein: (top inset) the vertical displacement from equilibrium of the upper leaflet u^+^ and (bottom inset) the slope normal to the protein. (B) Membrane upper and lower bounds of the mean hydrophobic core at the protein. Solid blue: the upper and lower mean leaflet locations; dashed red: 4^th^ order Fourier approximation. Dotted black: equilibrium leaflet positions. The boundary parameter is a measure of the distance around the boundary between the lipid-excluded surface of the protein and the membrane leaflet. (C) Solid and dotted blue: slopes of the membrane normal to the boundary. Dashed red: 4^th^ order Fourier slope approximations.

The boundary conditions for the membrane-protein contact curve were taken directly from the MD simulations. We used a 4^th^ order Fourier series (red dash) to fit the displacement along the boundary (blue) as shown in Figure *7*B for the parallel AH domain model. (See Fig. S2 for the 2L0J and 2N70 boundaries.) The pronounced pinching in the upper leaflet is most evident in this plot as can be seen by comparing the difference between the MD displacement curve and the equilibrium height +L_C_/2 (dashed curve). The slope along the inner boundary is much more noisy (solid and dotted blue curves) due to natural fluctuations as well difficulties in numerically taking derivatives of the mean membrane surfaces (Figure *7*C). The 4^th^ order Fourier series approximation (red dashed curves) smooths the MD data reducing this noise and avoids imposing high energy kinks on the boundary. Comparing panels B and C provides an idea of how membrane displacements are coupled to the membrane curvatures at the protein. Accurately solving the biharmonic equation requires knowledge of both of these features or knowledge of one and how it phenomenologically relates to the other. In our opinion, all-atom MD is currently the best approach for studying the coupling between both of these quantities. To our knowledge, the quantitative analysis shown here has never been examined before, and previous studies have either explicitly explored many different membrane slopes for a given displacement (Nielsen et al., 1998) or assumed (without justification) that the slope should be proportional to the membrane displacement from equilibrium (Choe et al., 2008).

Next, we numerically solved for the bilayer shape in a large, flat 200 Å × 200 Å patch using the inner boundary conditions taken from MD and the bilayer parameter values in Table 2. The far-field boundary conditions were clamped at zero slope and the upper and lower leaflets set to the equilibrium value L_C_/2 = 17.5 Å. A side-by-side comparison of the upper and lower leaflet heights from the MD (left column) and the continuum calculations (right column) for all three configurations shows that the model preserves many features present in the MD surfaces (Figure *8*). For instance, the upper leaflet shows significant curvature in the elastic calculations, while the lower leaflet is flat. The magnitude of the displacements is also similar in both cases, which is expected at the inner boundaries since they are identical along this curve. The spatial extent of the distortions between the MD and continuum model are similar, but the computed membrane height returns to the far field boundary value more quickly than the MD as seen for the upper leaflet of 2L0J in panel A. The MD relaxes over a distance of 10-17.5 Å while the continuum model does so over 7.5-12.5 Å. Another difference is that the continuum model produces much more smoothly varying solutions that show a higher degree of symmetry. The reduced noise in the surface is not surprising from this level of theory, but the increased symmetry was unexpected. For instance, 2L0J (panel A) shows a much higher degree of rotational symmetry (right) compared to the MD (left), and the upper leaflet distortion field for the parallel AH domain model (panel B) breaks into a clear 2-fold symmetric pattern with increased pinching along the X = -Y diagonal (right).

**Table 2.** Default elastic membrane material properties.

| Parameters | Values | Reference |
| --- | --- | --- |
| Membrane thickness ($L_c$) | 35 Å | this manuscript |
| Surface tension ($\alpha$) | $3.0 \times 10^{-13}$ N/Å | (Latorraca et al., 2014) |
| Bending modulus ($K_c$) | $L_c^2/24 \times K_a = 1.1 \times 10^{-9}$ NÅ | (Kim et al., 2012) |
| Gaussian modulus ($K_G$) | $-0.9 \times K_c$ | (Hu et al., 2012) |
| Areal compression modulus ($K_a$) | $2.13 \times 10^{-11}$ N/Å | (Rawicz et al., 2000) |

**Figure 8.**
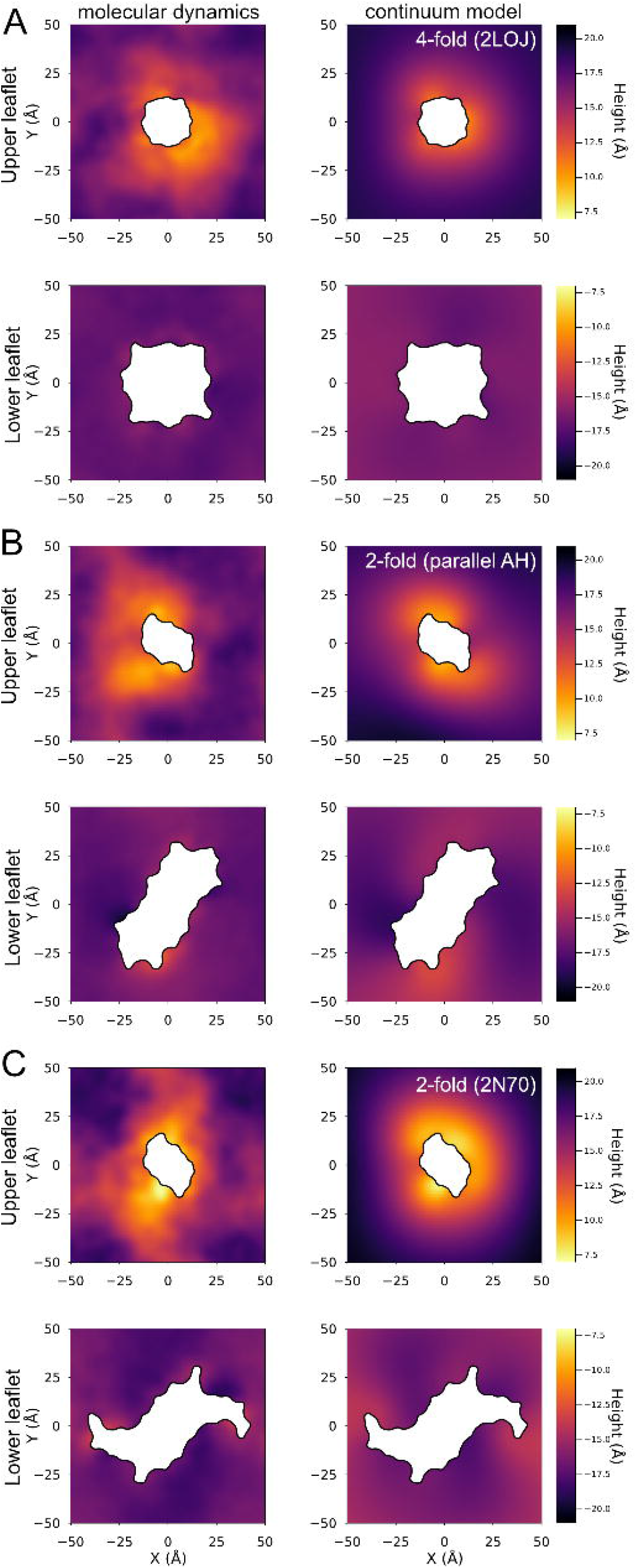
Continuum model membrane deformations compared to MD surfaces. Left column: MD upper and lower leaflet mean positions. Right column: continuum model minimum energy upper and lower leaflet surfaces for a flat membrane (200 Å by 200 Å membrane with zero displacement and slope on the outer boundary – entire patch not shown), with inner boundary conditions at the protein taken from the MD surfaces in the left column. (A) 2L0J. (B) Parallel AH domain model. (C) 2N70.

We believe that these differences are due to several factors including the elastic model penalizing higher order frequency modes and potentially insufficient averaging of the MD to remove low amplitude, high frequency features. However, the greatest difference between the MD and continuum solutions is likely that the MD has imposed periodic boundary conditions on a moderately sized membrane patch, while the continuum model has no imposed periodicity on a much greater region with flat, far-field boundaries twice as large in both linear dimensions.

We also predicted the total membrane deformation energy in a flat bilayer for each of these conformations. Given the significant bilayer compression and induced upper leaflet curvature near the protein boundary, we predict that all three conformations induce significant elastic deformation energy in the bilayer with the 4-fold 2L0J the lowest (32 kT), the 2-fold parallel AH domain model the middle (63 kT), and the largest is the 2-fold 2N70 structure (90 kT). While these values are substantial, they fall in the range of reported literature values for other computed membrane distortion energies induced by proteins such as 100 kT for lipid scrambling nhTMEM16 (Bethel & Grabe, 2016) and 50 kT for mechanosensitive gating of MscL (Ursell et al., 2007); the latter of which is consistent with the experimental total gating energy of 51 kT (Chiang et al., 2004). The membrane composition used here makes thick membranes (35 Å), and if we reduce the thickness to a value consistent with pure POPC bilayers (∼30 Å), our model predicts that the energy values drop by half to 14 kT, 31 kT, and 42 kT for 2L0J, parallel AH domain model, and 2N70, respectively (see Figure S3). While the 2-fold models induce higher membrane distortion energies, the total energy of the configuration is unknown since we have not quantified the energy of the other interactions in the system (protein enthalpy and entropy, protein-lipid interactions, protein-solvent interactions, etc.).

### C-2 symmetry broken channels prefer negative gaussian curvature

We used the continuum model to predict whether different M2 conformations would prefer different membrane geometries as assessed by the total membrane bending energy of different situations. To do this, we applied different far-field boundary conditions to impose global shapes on the membrane ranging from concave to convex spherical caps, which include flat membranes as the geometry transitions from concave down to concave up, as well as saddle geometries of pure negative Gaussian curvature. We assumed that the principal curvatures (κ_1_ and κ_2_) for each background shape were equal to each other (spherical caps) or opposite in sign (saddles), and we employed values corresponding to linear dimensions ranging from 100 Å radii of curvature (R_c_) to infinite in the case of the flat plane. Next, we assumed that each conformation retained its flat-bilayer boundary conditions as it migrates into regions of different curvature, thus we used the MD-generated boundary conditions (Figure *7*) on the inner boundary for all calculations. This assumption is based on the realization that failing to satisfy hydrophobic mismatch and lipid tilt at the membrane boundary can result in very large energy penalties, and the membrane adopts the same shape at the membrane-protein interface regardless of the global geometry. With these assumptions, we then computed the transfer free energy for each M2 configuration migrating from a region of flat curvature (i.e., the plasma membrane) into regions of different curvature, setting the reference membrane distortion in each case to zero for the flat reference case (Figure *9*). The X-axis in panel A of this figure, therefore, represents a continuum of shapes with NGC saddles on the left-hand side, flat bilayers at zero, and concave or convex caps on the right-hand side. Transfer free energies were then computed by calculating the difference between the flat empty bilayer plus the total membrane elastic energy of an M2 configuration embedded in a curved space minus the energy of the empty curved surface plus the M2 configuration embedded in a flat membrane (see Section S1 of Supporting Material). The flat empty bilayer in this cycle has no energy. This calculation is akin to a constant-area Helmholtz free energy in which the lipids displaced by the protein as it enters a curved space flow into the flat region evacuated by the protein, hence relieving all their stored strain energy.

**Figure 9.**
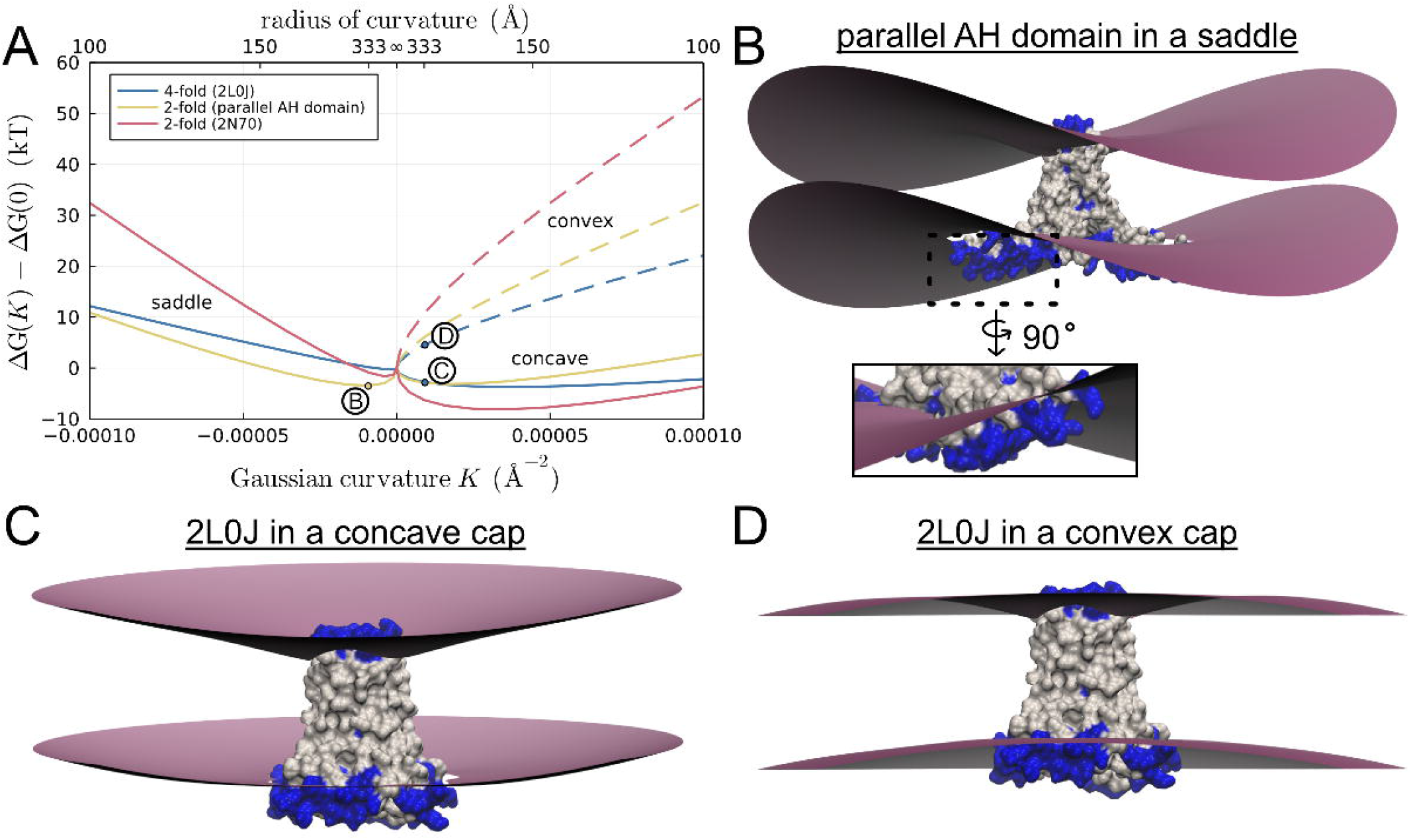
C2 symmetry broken conformations are stabilized in membranes with negative Gaussian curvature. (A) Transfer free energy ΔΔG(K) for moving M2 from a flat region to a region with Gaussian curvature K = ±1/R_c_^2^, where R_c_ is the radius of curvature (top X-axis). Solid lines at negative K show ΔΔG for three M2 models in saddles. Solid (dashed) lines at positive Gaussian curvature show ΔΔG in concave (convex) membranes. Labeled points on the plot indicate the energy for the membranes shown in panels B-D. For all shapes, κ_1_ = ±κ _2_. (B) Parallel AH domain model in a saddle. Inset: rotated close-up view of the lower leaflet distortion around the AH domain. (C) 2L0J in a concave spherical membrane. (D) 2L0J in a convex spherical membrane.

The C2-symmetry conformations are stabilized in saddles of NGC, and the parallel AH domain model is stabilized by 3 kT in a saddle with a 333 Å (33 nm) radius of curvature (gold curve at negative Gaussian curvatures in Figure *9*B). The figure and inset shows how the protein aligns in the membrane with its AH domains pointing along the concave principal curvature direction. It remains stable up to curvatures corresponding to ∼ 150 Å, which approaches the linear dimension of stalled viral buds. Contrarily, at this curvature, the 4-fold 2L0J is moderately destabilized in NGC by about 5 kT (blue curve at negative Gaussian curvatures).

All three M2 configurations preferentially sort to spherical caps of concave curvature (solid lines at positive Gaussian curvatures). Stabilization in this geometry is related to the destabilization in the convex caps – the pinching and membrane slope induced by the protein on the upper leaflet matches the global curvature of the concave caps (see Figure *9*C for 2L0J), hence reducing the total membrane deformation energy. This geometry is not relevant for a budding virus, but it does suggest that certain M2 channel configurations may become “trapped” in cellular regions rich in convex curvature such as caveolae. The calculations reveal that all three proteins are excluded by the convex spherical cap of a budding virus. At a radius of 125 Å corresponding to stalled buds observed in EM, the energy values range from 10 to 30 kT higher than the energy in a flat bilayer (dashed lines in Figure *9*A). These calculations corroborate the finding that M2 are not observed in the cap of stalled buds. Panel D shows the solution for the 4-fold 2L0J structure, and it is clear why the membrane bending energy is high; the pinching and membrane slope imposed by the protein on the upper (extracellular) leaflet is opposite to the natural curvature of the convex spherical cap. Since upper leaflet pinching is a feature of all M2 configurations studied here, they all are penalized in this geometry.

As noted earlier, membrane-protein interactions depend on the biophysical properties of the membrane, and membrane composition in the cell is extremely heterogeneous. To begin to address how the bilayer properties influence curvature sensing, we recomputed the energy values in Figure 9A in a thinner membrane with a 30 Å uncompressed hydrophobic thickness (Fig. S3). The general shapes of the curves are similar to those in Figure 9A; however, all of the energies are generally smaller in magnitude, as the membrane thickness is closer to the hydrophobic thickness of the protein and the relative amount of pinching in the upper leaflet is reduced. There are a few important quantitative differences in the curves, however. Most notably, proteins are significantly destabilized in thin concave curved membranes compared to thick membranes (Fig. S3).

## Discussion

Our simulations of the 4-fold symmetric 2L0J M2 channel reveal that the AH domains are highly dynamic and rearrange in the membrane, breaking both 4-fold and 2-fold symmetry of the *entire* channel, while the transmembrane portion experiences only minor deviations. Not only does the protein relax its conformation in the bilayer, but the membrane also relaxes its shape around the channel. Restrained simulations in three different M2 configurations exhibit common features in the induced membrane distortion patterns. We observed large curvature in the extracellular leaflets that was accompanied by pronounced upper-leaflet pinching, while the lower leaflets remained flat. Conversely, the lower leaflet lipids experienced greater tilt, while the upper leaflet lipids adopted near-bulk values. Simulation snapshots show that the increased tilt of the lower leaflet lipids is due to their tails “wrapping” around the partially inserted AH domains. Consequently, the increase in tilt decreases the vertical reach of the lower leaflet across the bilayer, and the upper leaflet pinches and bends to accommodate. Thus, our work suggests that the AH domains cause upper leaflet bending and compression regardless of how the individual AH domains are oriented with respect to the stable TM core. However, the simulations also indicate that the membrane distortion pattern adopts the symmetry of the channel; both 2-fold symmetric conformations induce a 2-fold symmetric deformation that has more upper-leaflet pinching and bending along one direction than another, while the 4-fold symmetric 2L0J simulation creates a more radially symmetric pattern.

Our continuum calculations provide an assessment of the membrane distortion energy induced by each conformation in a flat bilayer, and we computed substantial values on the order of 30-90 kT in the thick membranes (∼35 Å) simulated here, but more moderate values (14-42 kT) in thinner membranes (∼30 Å). In future work, we hope to verify these continuum predictions for the membrane energy using all-atom methodologies developed by the Faraldo-Gómez lab (Fiorin et al., 2020). Moreover, it would be informative to know the equilibrium probability of each of these three conformations, but this would require computing the total free energy of the entire system including the protein, which is beyond our current scope. Nonetheless, these systems allowed us to address an important question – do different channel conformations prefer different membrane geometries? The answer is yes, as our elasticity calculations in different background curvatures revealed that the membrane energy has a complex dependence on the magnitude of the curvature and surface shape. Qualitatively our results can be understood in terms of the MD generated distortion profiles in flat bilayers. The upper leaflet curvatures produced by all configurations favor migration into concave spherical regions and disfavor convex geometries, and the underlying mechanism is a relief of strain or increase in strain depending on the background geometry in a similar spirit to the curvature sensing model proposed by Johnson and co-workers (Fu et al., 2021). Thus, our work corroborates the finding that M2 is not enriched in the convex spherical cap of budding virions. Both C2 symmetric conformations favor saddle regions of moderate NGC, while the 4-fold 2L0J does not. Within the plane of the membrane the 2-fold symmetric proteins have a long and a short axis, and they align with the principal curvatures of the saddle to minimize the membrane elastic energy as shown in Figure *9*B. However, the parallel AH domain model is ∼3.5 kT more stable in the saddle relative to a flat bilayer, while the 2N70 structure is only about ∼2 kT. While these energy differences are modest, they could lead to enrichments in saddles over the flat regions of the plasma membrane by 10-fold or more, and their presence would likely stabilize saddle formation.

There are a variety of M2 structures. We call attention to two distinguishing properties of these structures: the helicities of the amphipathic domains and the degrees of symmetry. 2L0J may represent M2 in a membrane environment without curvature, whereas our work follows others (Elkins et al., 2017) in supposing 2N70 may represent M2 in a highly curved environment, one possibly more curved than the physiological case. Indeed, many differences in M2 structures could be partially accounted for by considering the curvature of the membrane mimetic. C2 symmetry of the membrane deformation pattern appears energetically significant to M2’s negative curvature sensitivity. Helicity in the amphipathic domain does not appear as significant, but it may contribute to structural changes necessary to access the C2 conformations competent for budding.

Compared to the CD constructs considered here, full length M2 has an extended C-terminal domain that is disordered. While the structural implications are far from certain, it is likely that the full-length protein is even more dynamic in the entire C-terminal region to accommodate conformational entropy for the disordered tail. DEER studies (Herneisen et al., 2017) also suggest that full length M2 may have a greater propensity to explore C2 conformations than the CD constructs. The dynamics observed here were already sufficient to complicate analysis of membrane deformation patterns around M2, so proper interpretation of simulation studies of full-length M2 presents a challenge for future work.

Understanding the role of M2 in viral egress could lead to novel therapeutic strategies not just for influenza, but other viruses or pathogens that bend mammalian plasma membranes. Most M2 drugs have historically been pore blockers, but drugs antagonizing C2 tetramerization at an external site may merit additional exploration. To our knowledge, no drugs are presently known to specifically inhibit M2 NGC binding and subsequent viral release. Early M2 structures showed rimantadine binding at an external site, but it was later determined to be a lower affinity site compared to the pore. Finding higher affinity ligands for this alternate rimantadine site may yield additional insight into the drugability of M2’s membrane bending function. Additionally, cholesterol has been shown to bind at two of the tetramer interfaces (Elkins et al., 2017), suggesting that specific binding to this shallow groove is possible. However, it is not clear whether existing structural information is sufficient to find binders at the peripheral site: 2N70 likely represents an extreme C2 conformation, not the C2 conformation present on average at the budding neck. Further studies combining unrestrained MD with continuum energy calculations may be able to find dynamically accessible conformations with low energy in saddle-shaped membranes, and such conformations could be fruitful poses for docking or drug design, thus opening up the possibility for using small molecules to stabilize M2 channels in fission-incompetent conformations.

## Methods

### Molecular Dynamics

#### Setup

Simulations of the M2 conductance domain were initiated from PDBID 2L0J (residues 22-62) (Sharma et al., 2010b), the parallel AH domain model based on PDBID 2N70 (residues 22-60), and PDBID 2N70 (residues 18-60) (Andreas et al., 2015). Residues were assigned standard protonation states at pH 7, with the exception of His37 which retained the protonation states from PDBID 2L0J (doubly protonated on chains A and C, epsilon protonated on chains B and D). Structures were embedded in 4:1 POPC:POPG bilayers, with cholesterol-containing simulations containing 30 % mole fraction cholesterol (56:14:30 POPC:POPG:cholesterol), and solvated with 150 mM KCl resulting in a neutralized system using CHARMM-GUI (Lee et al., 2016). The force fields used for protein, lipids, and water were CHARMM36m (Huang et al., 2017), CHARMM36 (Klauda et al., 2010), and TIP3P (Jorgensen et al., 1983), respectively. Standard CHARMM parameters were used for ions (Vanommeslaeghe et al., 2010).

#### Production

Simulations were run on the Wynton HPC cluster with GPU nodes using GROMACS 2018 (Abraham et al., 2015). All simulations were minimized and equilibrated using the default options provided by CHARMM-GUI, excepting fully restrained simulations which retained backbone restraints with a force constant of 4000 kJ/mole/nm^2^ throughout. Briefly, minimization was performed for 5000 steps with protein backbone, protein side-chain, and lipid harmonic positional restraints at force constants of 4000, 2000, and 1000 kJ/mole/nm^2^, respectively, as well as dihedral restraints with a force constant of 1000 kJ/mole/nm^2^. A multi-step equilibration protocol stepped down the restraints over a series of 2 ns phases based on recommended CHARMM-GUI defaults. Following equilibration, an unrestrained series of pre-production simulations totaling 10 ns was run using a 2 fs time step, a Parrinello-Rahman barostat (Parrinello & Rahman, 1981) with semi-isotropic pressure control at 1 atm, and a Nose-Hoover thermostat (Hoover, 1985; Nose, 1984) set to 303.15 K. Nonbonded interactions were cut off at 12 Å with force-switching between 10 and 12 Å, long-range electrostatics were calculated with particle mesh Ewald (Darden et al., 1993), and hydrogens were constrained with the LINCS algorithm (Hess et al., 1997). Final production MD proceeded using the same options with hydrogen mass repartitioning (Balusek et al., 2019) enabled to allow for use of a 4 fs timestep.

#### Analysis

Systems were first centered on the protein and wrapped to make molecules whole, then rotated and translated (constrained to the XY-plane) to maintain the starting configuration using GROMACS (gmx trjconv). Membrane surface calculations were performed using a custom analysis package based on MDAnalysis (Michaud-Agrawal et al., 2011), NumPy (Harris et al., 2020), and SciPy (Virtanen et al., 2020) with the same approach as outlined previously (Bethel & Grabe, 2016). Briefly, we erect a rectilinear grid with 1 Å spacing everywhere except at the protein-membrane interface, where we use a level set method based on the protein structure to move adjacent membrane grid points onto the surface (Argudo et al., 2017). Then, the positions of C22 and C32 atoms on POPC and POPG residues (first carbon atoms of each lipid tail) from MD simulations were interpolated onto this distorted grid using SciPy’s implementation of the Clough-Tocher scheme to construct, for every time point analyzed, a hydrophobic surface for each leaflet. To avoid edge effects, the simulation frames were expanded by mirroring lipids within 18 Å of the box edge, producing an expanded box of 126 Å × 126 Å over which grids were interpolated and then trimmed back down to the true box size. This produces interpolated grids that vary smoothly over the entire 95 Å × 95 Å grid and capture the full periodicity of the system. Bilayer hydrophobic thicknesses were then calculated by taking the difference between interpolated upper and lower leaflet hydrophobic surfaces. All surfaces were then averaged across timepoints. These profiles were visualized using MATLAB R2015R [MathWorks, Natick, MA, USA], Matplotlib (Hunter, 2007), Seaborn (Waskom, 2021), Plots.jl (Christ et al., 2022), and Julia (Bezanson et al., 2012).

Lipid tilt angles were calculated per simulation frame for every phospholipid. The tilt angle was defined as the angle between the Z axis of the box, taken as the bilayer normal, and the vector pointing from the phosphorous atom of each lipid phosphate group to the midpoint between the last carbon atoms of each lipid tail. We interpolated a tilt surface using the same grid and method as for the membrane surfaces just described. Tilt surfaces for each timepoint are then averaged together across the full production time. Average membrane properties were computed over the second half of each trajectory. See Table 1 for a list of simulation times.

### Continuum Membrane Mechanics Model

Continuum elastic numerical calculations were performed on 200 Å × 200 Å bilayer patches, which are sufficiently large to allow local distortions at the protein to relax (decay length λ_D_ = (1/γ)^½^ ∼ 60 Å, where γ = α/K_C_), while being small enough to capture curvatures relevant to viral budding. Smaller patches induced higher energetic penalties, while increasing the patch size did not significantly reduce the elastic energy. The protein-membrane boundary positions were taken from the intersections of the mean MD membrane surfaces with the lipid-excluded surface of the proteins (i.e., the molecular surfaces using a 4.58 Å probe radius). The inner slope boundary conditions were set to the normal component of the gradient of the MD mean surfaces. Outer boundary conditions were set to impose a flat bilayer, a spherical cap, or saddle background. With and without protein inclusions, for spherical geometry, the outer boundary upper *u*^+^ and lower *u*^-^ leaflet displacements were set to:

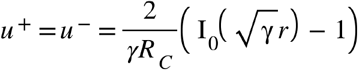

while for saddles, the displacements were set to:

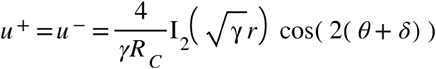

where I_0_ and I_2_ are modified Bessel functions of the first kind, r and θ are polar coordinates, and δ is an arbitrary constant (see Fig. S4 and Section S2.3 for derivation.) The outer boundary slopes were set to the boundary-normal component of the gradients of the above expression. In an empty membrane, with the above boundary conditions, these Bessel function solutions are solutions of the Euler-Lagrange equations throughout the membrane, and our numerical solver produced surfaces of these forms. The small *r* limiting forms indicate that near the origin, these boundary conditions produce surfaces with Gaussian curvatures of K = ±1/R_c_^2^.

We calculated the difference in minimum energy configurations between membranes with and without M2 for surfaces with a range of Gaussian curvatures. For simplicity, we only considered surfaces whose principal curvatures K_1_ and K_2_ were the same magnitude: |κ_1_| = |κ_2_| = 1/R_c_. The only free parameters were a constant z_0_ offset and the orientation of the saddle δ. For spherical caps, z_0_ was optimized using Brent’s method. For saddles, z_0_ and δ were optimized using the Nelder-Mead method. Searches were implemented in Julia using Optim (Mogensen & Riseth, 2018).

## Acknowledgements

We would like to thank Bill DeGrado, Yongcheng Zhou, Kathleen Howard, Matthew Jacobson, and members of the Grabe lab for helpful discussions regarding this work. This work was supported by funds from the UCSF Biophysics Program, NIH postdoctoral fellowship to J.L. (4T32HL007731-28), and NIH grants R01GM117593 and R01GM137109. Hardware for simulations was provided by NIH R01GM089740.

