## Supporting Material for "Membrane curvature sensing and symmetry breaking of the M2 proton channel from Influenza A"

### S1. Membrane elastic energy of M2 in curved membranes

The geometric shape and hydrophobic mismatch between a transmembrane protein and the surrounding lipid bilayer can distort the shape of the membrane and change the total membrane energy. The protein-induced membrane deformation energy may be lower in some curved membrane regions than others, leading to curvature sensing. Using our numerical continuum elasticity solver[1], we explored which background membrane curvatures are energetically favorable to M2, based on the deformations of lipid bilayers around M2 observed in molecular dynamics (MD) simulations.

Let $G_p(\kappa_1, \kappa_2)$ be the membrane elastic energy around a protein in a membrane region with principal curvatures $\kappa_1$ and $\kappa_2$, and compare this to the membrane elastic energy $G_e(\kappa_1, \kappa_2)$ of a similarly curved membrane region without protein inclusions:

$$\Delta G(\kappa_1, \kappa_2) = G_p(\kappa_1, \kappa_2) - G_e(\kappa_1, \kappa_2). \tag{S1}$$

We want to compare the protein-induced membrane energy difference in a curved region to that in a flat region. Let $\Delta\Delta G(\kappa_1, \kappa_2) = \Delta G(\kappa_1, \kappa_2) - \Delta G(0, 0)$:

$$\begin{aligned} \Delta\Delta G(\kappa_1, \kappa_2) &\coloneqq \Delta G(\kappa_1, \kappa_2) - \Delta G(0, 0) \\ &= (G_p(\kappa_1, \kappa_2) - G_e(\kappa_1, \kappa_2)) - (G_p(0, 0) - G_e(0, 0)) \\ &= (G_p(\kappa_1, \kappa_2) + G_e(0, 0)) - (G_p(0, 0) + G_e(\kappa_1, \kappa_2)). \end{aligned} \tag{S2}$$

That is, $\Delta\Delta G(\kappa_1, \kappa_2)$ is the change in membrane energy when a protein is moved from flat region to a curved region, while simultaneously displacing lipids from the curved region to the flat region. Elsewhere for brevity we will refer to $\Delta\Delta G(\kappa_1, \kappa_2)$ as $\Delta\Delta G(K)$, where $K = \kappa_1\kappa_2$.

### S2. Minimum energy surfaces

To set up numerical calculations in regions of prescribed curvature, we found analytic minimum energy solutions of known curvature, and used them as boundary conditions for our numerical solver. In a protein-free region, the Euler–Lagrange equations for the vertical displacements from equilibrium of the upper and lower membrane leaflets, $u^+$ and $u^-$ respectively, of an intrinsically flat membrane bilayer are

$$\nabla^4 u^+ - \gamma\nabla^2 u^+ + \beta(u^+ - u^-) = 0, \text{ and} \tag{S3}$$

$$\nabla^4 u^- - \gamma\nabla^2 u^- + \beta(u^- - u^+) = 0, \tag{S4}$$

where $\gamma = \alpha/K_c$ is the ratio of the stretch modulus $\alpha$ to bending modulus $K_c$, and $\beta = 2K_a/K_c/L_c^2$ is the ratio of the compression modulus $K_a$ scaled by the membrane thickness $L_c$ squared to the bending modulus. Solving these forth order differential equations requires specifying two sets of boundary conditions for each leaflet. We chose to specify $u^+$ and $u^-$, and their normal derivatives $\nabla u^+ \cdot \hat{n}$ and $\nabla u^- \cdot \hat{n}$, where $\hat{n}$ is the direction normal to the boundary.

Solutions to these equations are well know. For example, Nielsen[2] solved the case of equal magnitude but opposite sign displacements and slopes. We instead looked at solutions with identical offsets and slopes on the boundaries. When $u^+ = u^- = u$, and $\nabla u^+ \cdot \hat{n} = \nabla u^- \cdot \hat{n} = \nabla u \cdot \hat{n}$ everywhere on the boundary, the two leaflets decouple, and minimal surfaces are solutions of the homogeneous equation

$$\nabla^4 u - \gamma \nabla^2 u = 0. \tag{S5}$$

In polar coordinates, the solution to this simpler biharmonic equation is

$$u(r, \phi) = C + D\ln(r) + \sum_{m=0}^{\infty} \left[ A_m \mathrm{I}_m(\sqrt{\gamma} r) \cos(m(\phi + \delta_m^{\mathrm{I}})) + B_m \mathrm{K}_m(\sqrt{\gamma} r) \cos(m(\phi + \delta_m^{\mathrm{K}})) \right], \tag{S6}$$

where $\mathrm{I}_m$ and $\mathrm{K}_m$ are order $m$ modified Bessel functions of the first and second kind, and $C$, $D$, $A_m$, $B_m$, $\delta_m^{\mathrm{I}}$, and $\delta_m^{\mathrm{K}}$ are constants determined by the boundary conditions. We were interested in exploring solutions which are finite at the origin and look either spherical ($m = 0$) or saddle–like ($m = 2$).

#### S2.1. Spherical caps

A hemispherical cap touching the origin, centered at $z = R$ above the origin, has height $z_{\mathrm{cap}}$:

$$\begin{aligned} z_{\mathrm{cap}}(r) &= R - \sqrt{R^2 - r^2} \qquad \text{(S7)} \\ &= \frac{r^2}{2R}\left(1 + \frac{r^2}{2R^2} + \mathcal{O}(\frac{r^4}{R^4})\right) \\ &\approx \frac{r^2}{2R} \text{ for } r \ll R. \end{aligned}$$

The Maclaurin series for $\mathrm{I}_0(\sqrt{\gamma} r)$ is

$$\mathrm{I}_0(\sqrt{\gamma} r) = 1 + \frac{\gamma r^2}{4} + \frac{\gamma^2 r^4}{64} + \mathcal{O}((\sqrt{\gamma} r)^6), \tag{S8}$$

so an empty membrane patch which looks locally like a spherical cap with radius of curvature $R$ is given by the $m = 0$, rotationally symmetric membrane with solution

$$\begin{aligned} u_{\mathrm{cap}}(r, \phi) &= \frac{2}{\gamma R}\left(\mathrm{I}_0(\sqrt{\gamma} r) - 1\right) \qquad \text{(S9)} \\ &\approx \frac{r^2}{2R} \text{ for } \gamma r^2 \lesssim 1. \end{aligned}$$

With $\alpha = 3 \times 10^{-13}$ N/Å and $K_c = 1.1 \times 10^{-9}$ NÅ, $\gamma = \alpha/K_c = 2.7 \times 10^{-4}$ Å$^{-2}$. The membrane is expected to deviate from a spherical cap when $r \sim \sqrt{1/\gamma} = 60.2$ Å. Fig. S4A shows the ideal mathematical spherical cap and the biharmonic solution used to simulate a spherical cap.

#### S2.2. Saddles

One definition of a unit saddle is $z = (x^2 - y^2) = \frac{1}{2}r^2\cos(2\phi)$. This saddle has principal curvatures $\kappa_1 = -\kappa_2 = 1$. To match our solutions above, we add in an arbitrary rotation by angle $\delta$, and scale the principal curvatures by $\kappa_{1/2} = \pm 1/R$:

$$z_{\text{saddle}} = \frac{r^2}{2R}\cos(2(\phi+\delta)), \tag{S10}$$

where as $R \to \infty$ the surface becomes flat. The $m = 2$ saddle-like membrane deformation from equation S6 is

$$\begin{aligned} u_{\text{saddle}}(r,\phi) &= A_2\,\mathrm{I}_2(\sqrt{\gamma}r)\,\cos(2(\phi+\delta_2^{\mathrm{I}})) \qquad \text{(S11)}\\ &= A_2\left[\frac{\gamma r^2}{8} + \frac{\gamma^2 r^4}{96} + \mathcal{O}((\sqrt{\gamma}r)^6)\right]\cos(2(\phi+\delta_2^{\mathrm{I}})) \end{aligned}$$

Setting $A_2 = \frac{4}{\gamma R}$ and $\delta_2^{\mathrm{I}} = \delta$ gives a minimum energy saddle–like surface such that $u_{\text{saddle}} \approx z_{\text{saddle}}$ when $\gamma r^2 \lesssim 1$. Fig. S4B shows the ideal mathematical saddle and the biharmonic solution used to simulate a saddle.

#### S2.3. External boundary conditions

For the boundary conditions along the protein-membrane contact curve, we used the values derived from MD simulations (Figs. 7, S2), and assumed that these boundary conditions do not significantly change as the curvature of the surrounding membrane patch changes, since deviations of the local boundary conditions will result in high energy penalties in the hydrophobic, electrostatic, and steric energies.

For spherical caps of radius $R$, the external boundary conditions were set to

$$u^+ = u^- = u_{\text{cap}} = \frac{2}{\gamma R}\left(\mathrm{I}_0(\sqrt{\gamma}r) - 1\right), \text{ and} \tag{S12}$$

$$\nabla u^+ \cdot \hat{n} = \nabla u^- \cdot \hat{n} = \nabla u_{\text{cap}} \cdot \hat{n} = \frac{2}{\sqrt{\gamma}R}\mathrm{I}_1(\sqrt{\gamma}r)\,\hat{r}\cdot\hat{n} \tag{S13}$$

The sign of the radius of curvature determines if the cap is convex or concave from the extracellular side. For saddles with principal curvatures $1/R$ and $-1/R$, the external boundary conditions were set to

$$u^+ = u^- = u_{\text{saddle}} = \frac{4}{\gamma R}\mathrm{I}_2(\sqrt{\gamma}r)\,\cos(2(\phi+\delta)), \text{ and} \tag{S14}$$

$$\begin{aligned} \nabla u^+ \cdot \hat{n} = \nabla u^- \cdot \hat{n} &= \nabla u_{\text{saddle}} \cdot \hat{n} \\ &= \frac{2}{\sqrt{\gamma}R}\left[\mathrm{I}_1(\sqrt{\gamma}r) + \mathrm{I}_3(\sqrt{\gamma}r)\right]\cos(2(\phi+\delta))\,\hat{r}\cdot\hat{n} \\ &\quad - \frac{8}{\gamma r R}\mathrm{I}_2(\sqrt{\gamma}r)\,\sin(2(\phi+\delta))\,\hat{\phi}\cdot\hat{n}. \qquad \text{(S15)} \end{aligned}$$

### S3. Elastic energy in a thinner membrane

The mean hydrophobic core thickness far from the protein in the three MD simulations, based on the POPC and POPG C22 and C23 atoms, was 35 Å. Therefore we used 35 Å for the continuum membrane solver. Here we reproduce the energetic profiles in Fig. 9, but for a membrane with core thickness $L_c = 30$ Å. As in the main text, we set the membrane bending modulus $K_c = L_c^2/24 \times K_a$. The membrane energy for the thinner membrane was then minimized again over the free parameters ($z_0$ and $\delta$).

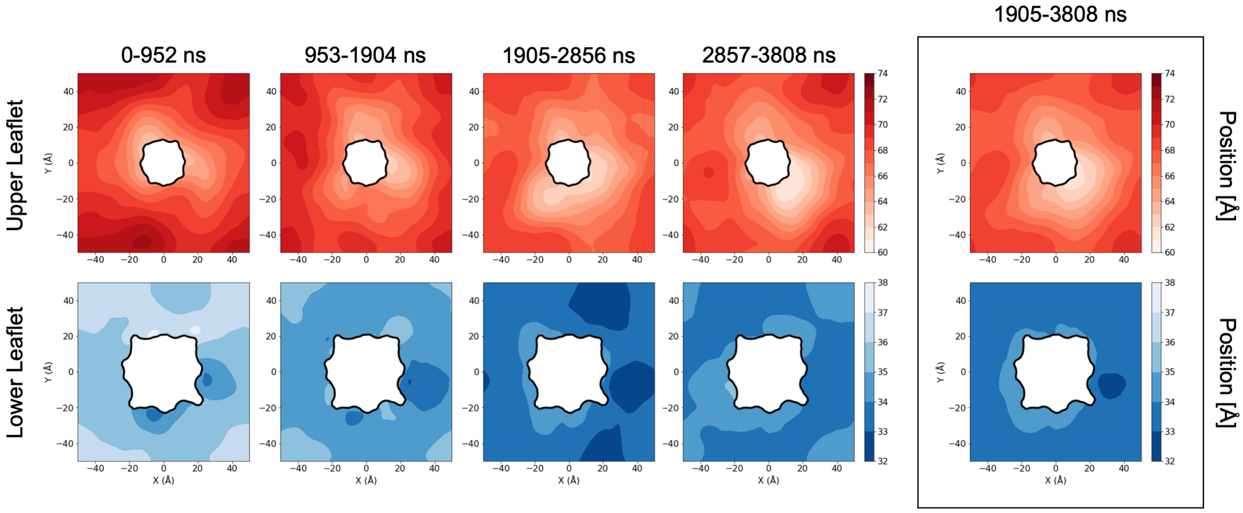


Figure S1: Equilibration of leaflet heights in the restrained 2L0J simulation. The four columns on the left show block-averaged heights (see color bars) for the upper (top row) and lower (bottom row) leaflets across each quarter of the full-length simulation, while the boxed panels at the right show the leaflet surfaces used as production data for analysis and discussion in the manuscript, made by averaging across the second half of the simulation. Both leaflets relax downward by $\sim$ 4-5 Å across the first half of the simulation and then show reduced fluctuations, indicative of sampling around equilibrium, in the second half. Contour levels for each leaflet are spaced 1 Å apart.

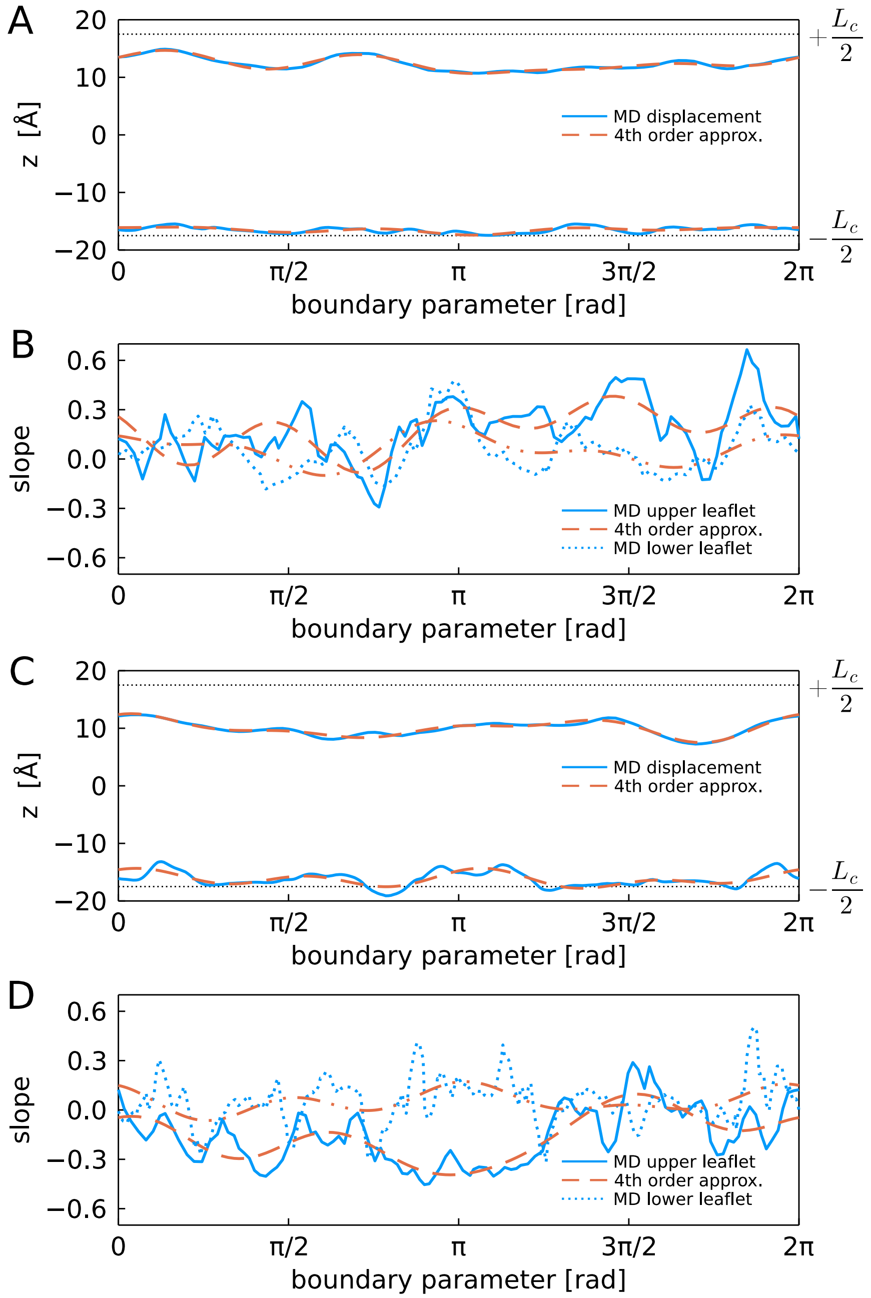


Figure S2: Boundary conditions extracted from MD simulations. A,B: 4-fold (2L0J). C,D: 2-fold (2N70). A,C: Upper and lower bounds of the mean hydrophobic core at the protein. Solid blue: mean MD membrane hydrophobic boundaries. Dashed red: Forth order Fourier series approximation of the MD boundary. B,D: Slopes of the mean MD hydrophobic surfaces at the protein in the direction normal to the boundary. Blue solid (dotted): upper (lower) leaflet slopes. Dashed red: Forth order Fourier series approximation of the MD slopes. The boundary parameter is the length around the boundary, normalized to $2\pi$.

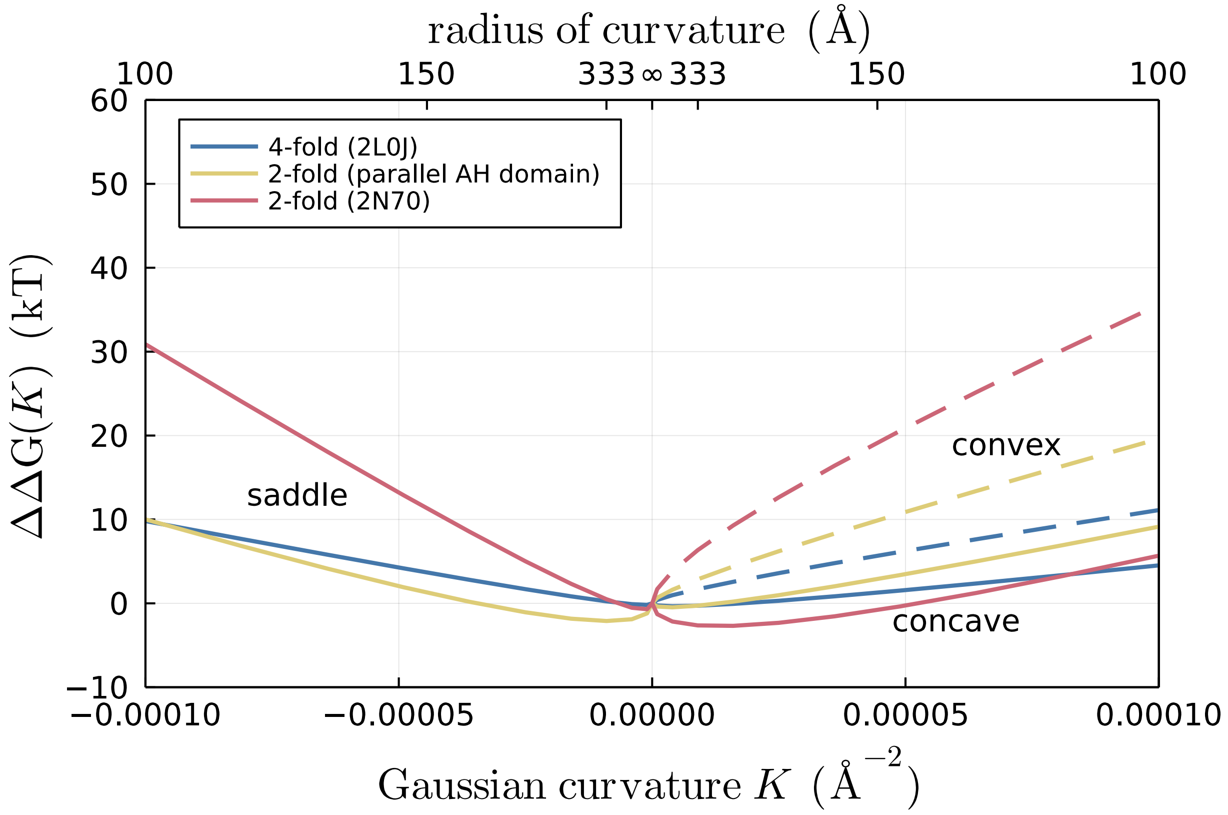


Figure S3: Influence of a thin membrane on curvature sensing by different M2 channel conformations. Here, a thinner membrane (30 Å uncompressed hydrophobic core) was used to compute the total membrane bending energy for moving from a flat membrane to different curvature regions compared to the 35 Å uncompressed hydrophobic length employed in Fig. 9A. Positive Gaussian curvatures $K$ correspond to concave (solid curves) or convex (dashed curves) spherical caps, while negative Gaussian curvature values correspond to saddles.

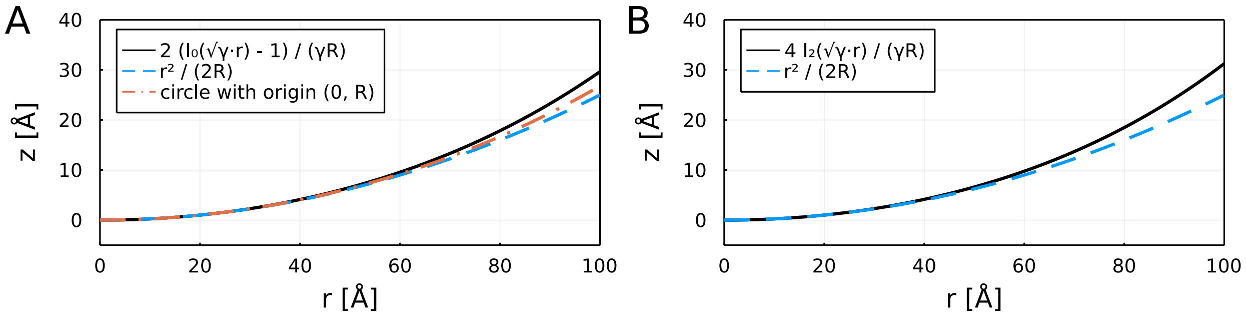


Figure S4: Membrane shapes which approximate an ideal spherical cap or saddle. A. Dash–dotted red: circle of radius R = 200 Å. Dashed blue: quadratic approximation of circle, valid for $r \ll R$. Solid black: the m = 0 analytic biharmonic solution finite at the origin behaves quadratically for $\sqrt{\gamma} r \lesssim 1$. B. Dashed blue: the ideal saddle $z = \cos(2\phi)\, r^2/(2R)$ is quadratic along the $x$ and $y$ axes with radii of curvature $R$ and $-R$, respectively. Solid black: the m = 2 analytic biharmonic solution diverges from quadratic when $r \gtrsim \sqrt{1/\gamma} \approx 60.2$ Å. The numeric solutions from the membrane solver are not shown because they are indistinguishable from the analytic solutions at this scale.
